# SATB2 organizes the 3D genome architecture of cognition in cortical neurons

**DOI:** 10.1101/2022.12.27.522010

**Authors:** Nico Wahl, Sergio Espeso-Gil, Paola Chietera, Aodán Laighneach, Derek W. Morris, Prashanth Rajarajan, Schahram Akbarian, Georg Dechant, Galina Apostolova

## Abstract

*SATB2* is genetically associated with human intelligence. Since SATB2 protein structure predicts a function in DNA looping, we analyzed the impact of SATB2 on 3D genome architecture and chromatin accessibility in cortical neurons. Our data reveal strong effects of SATB2 on chromatin looping between enhancers and promoters of neuronal activity-regulated genes, which closely correlate with gene expression. We furthermore identify SATB2-dependent alterations at all 3D genome architectural levels, including compartments, Topologically Associated Domains and Frequently Interacting Regions. Genes linked to SATB2-dependent 3D genome changes are implicated in highly specialized neuronal functions and contribute to cognitive ability and risk for neuropsychiatric and neurodevelopmental disorders. The altered non-coding regions are enriched for common variants associated with educational attainment, intelligence and schizophrenia. Our data establish SATB2 as a 3D genome organizer, which operates both independently and in cooperation with CTCF to set up the chromatin landscape of pyramidal neurons for cognitive processes.

## Introduction

Cognitive processes require a dynamic functional link between synaptic activity and gene expression in the neuronal nucleus^1^. Emerging evidence suggests a role of spatial genome organization in both normal cognition and cognitive impairment^2^. Recent studies on a small number of candidate genes indicate that organization of DNA in chromatin loops contributes to genetic risk architecture of cognitive diseases with early childhood/young adulthood onset, including autism spectrum disorder (ASD) and schizophrenia (SZ)^3,4^. It has also been observed that structural DNA variants associated with neuropsychiatric disorders are positioned in intergenic or intragenic non-coding sequences and influence neuronal gene expression by bypassing the linear genome to directly interact with target genes^5,6^. However, specialized molecular mechanisms that affect the higher-order chromatin landscape of neuronal gene regulatory networks remain poorly understood. Here, we address this question by studying the function of SATB2, a nuclear protein that in the CNS is selectively expressed in pyramidal neurons of cortex and hippocampus^7–9^, two brain regions inextricably connected to cognitive function.

A growing body of evidence has linked SATB2 to higher brain functions. Genome-wide association studies (GWAS) have identified *SATB2* as a locus associated with human cognitive ability^10^. Common variation in the genes regulated by SATB2 or encoding SATB2-interacting proteins influences human cognitive ability and contribute to SZ^11,12^. Rare *de novo* mutations within *SATB2* locus cause SATB2-associated syndrome (SAS), an autosomal dominant disorder characterized by developmental delay and severe intellectual disability^13^. Mouse mutants, in which SATB2 is selectively deleted in adult forebrain pyramidal neurons, display cognitive defects, including impaired late LTP and long-term memory dysfunction^14,15^. Potent effects of SATB2 on neuronal gene transcription include regulation of activity-dependent immediate early genes (IEGs)^16^. Consistent with a role in neuronal plasticity, SATB2 protein levels are regulated by BDNF and synaptic activity^14^.

The protein structure of SATB2 predicts a function as a chromatin loop organizer. SATB2 binds to DNA via four different DNA-binding domains: a homeodomain, two CUT domains, and a CUT-like domain^17,18^. Self-association of SATB2 into homomeric di-or tetramers is mediated by an ubiquitin-like domain at the N-terminus^18,19^. SATB2-homomeric complexes, composed by monomers bound to distant genomic regions, might promote formation and/or stabilization of intra-chromosomal loops. While recent studies in non-neuronal cell types have provided evidence for alterations in chromatin accessibility and individual chromatin loops upon SATB2 loss a genome-wide analysis of SATB2 effects on 3D-chromatin structure is currently lacking^20–22^.

In this study, we identify epigenomic and higher-order chromatin interactions that depend on SATB2 in cortical pyramidal neurons, by integrating high-resolution, multidimensional datasets from *Satb2* conditional knockout (cKO) and floxed mice. We observe robust and highly coordinated alterations in both chromatin accessibility and chromatin loop landscape upon SATB2 deletion. Surprisingly, we find that SATB2 effects on 3D-genome organization are not limited to modifying chromatin loops but also encompass effects on large-scale architectural levels, including A/B compartments, Topologically Associated Domains (TAD) and Frequently Interacting Regions (FIRE). With remarkable specificity, SATB2-dependent 3D-epigenome remodeling occurs at genomic loci that are functionally associated with cognition and synaptic signaling, and contribute to human cognitive ability and risk for neuropsychiatric and neurodevelopmental disorders.

## Results

### SATB2 deletion in cortical neurons causes extensive alterations in chromatin accessibility and chromatin loop landscape

To explore a role of SATB2 in shaping 3D genome architecture, we compared chromatin interactions and chromatin accessibility upon SATB2 ablation by using cultured cortical neurons derived from *Satb2*^flx/flx^::*Nes-Cr*e (*Satb2* cKO) and *Satb2*^flx/flx^ (floxed) mice^12^ (Fig. S1A). While *ex vivo* collected neurons often vary in their activity state, ranging from silent to highly active, cell-to-cell heterogeneity in our primary cultures was kept at minimum by addition of bicuculline, a GABA-A receptor antagonist, which stabilizes moderate neuronal activity in all neurons^23^. *Satb2* cKO and floxed cultures consisted of more than 85 % pyramidal neurons, did not differ morphologically, and expressed developmental and synaptic maturation markers at equal levels (Fig. S1B). To identify changes in chromatin accessibility and chromatin interactions, three ATAC-seq and six Hi-C libraries were prepared per genotype, respectively. To map SATB2 binding sites genome-wide, two CUT&RUN libraries were prepared from floxed cultures^24^. ATAC-seq, CUT&RUN, and Hi-C libraries were highly reproducible across replicates^25^ (Fig. S1C-D). Furthermore, Hi-C contact maps were highly similar to a previously reported *in vivo* Hi-C dataset derived from NeuN^+^ sorted neurons of adult mouse cortex^26^ (Fig. S1D).

The analysis of SATB2 genome-wide occupancy by CUT&RUN yielded a total of 14964 binding sites. Similar to previous observations in hippocampal neurons^14^, we found a preferential binding of SATB2 to promoters also in cortical neurons, with 7819 peaks overlapping with promoters (Fig. 1A-B). Chromatin state modeling by ChromHMM using neonatal mouse forebrain datasets^27,28^ further revealed that SATB2 peaks are associated with active chromatin states, including several types of enhancers (Fig. 1C). We next performed a Gene Ontology (GO) analysis to identify enriched pathways within the genes harboring SATB2 binding sites at their promoters. We observed a strong overrepresentation of genes involved in synapse organization, regulation of membrane potential, and ion transmembrane transport, such as *Cacna1c, Grin2a, Shank1* and *Magi2* (Fig. 1D).

**Figure 1:**
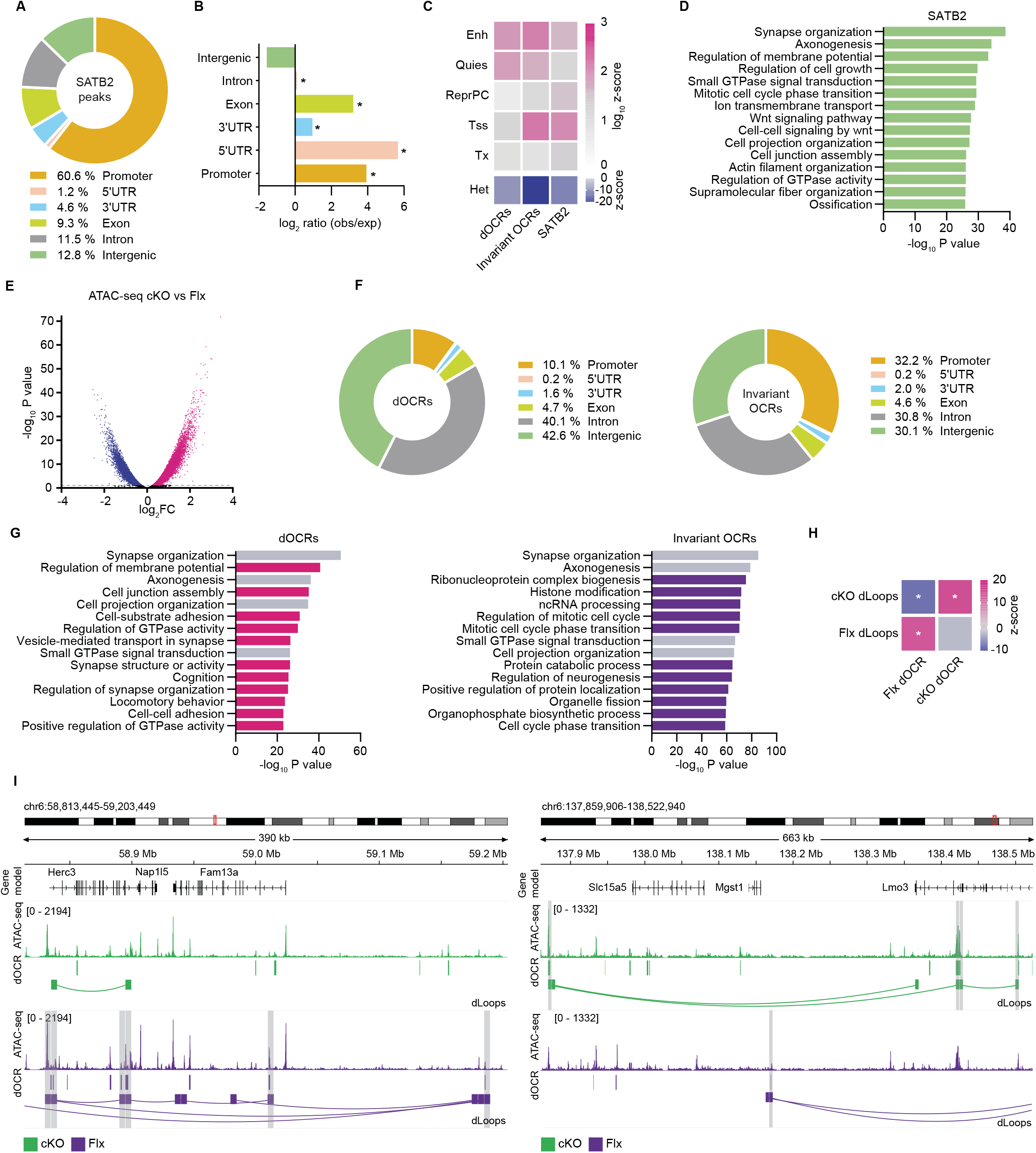
3D epigenome remodeling upon SATB2 deletion in cortical neurons. A. Genomic distribution of SATB2 binding sites (n = 2). Pie chart showing the percentage of regulatory elements overlapping with SATB2 peaks. B. Enriched genomic annotations for SATB2 peaks. P values were calculated by HOMER tools (*P < 0.001). C. Enrichment analysis of dOCRs, invariant OCRs, and SATB2 peaks over pooled chromatin states (see Methods), inferred by P0 forebrain ChromHMM 18-state model, for all enrichments and depletions, P < 0.00001. D. Gene Ontology (GO) analysis for biological processes of gene promoters occupied by SATB2. E. Volcano plot illustrating differential chromatin accessibility between cKO and floxed cortical neurons (FDR < 0.01, n = 3). F. Genome-wide distribution of dOCRs and invariant OCRs, represented as a percentage. G. GO analysis of genes assigned to dOCRs and invariant OCRs. Peak annotation is based on seq2gene function of ChIPseeker^29^. GO categories, specific to dOCRs and invariant OCRs, are highlighted in pink and purple, respectively. H. Enrichment of Flx dOCRs at the anchors of Flx dLoops, and vice versa, of cKO dOCRs at the anchors of cKO dLoops. Color bar shows the z-score, as calculated by permutation tests with regioneR, *P < 0.00001. I. Representative IGV genome browser tracks illustrating the overlap between Flx dOCRs and Flx dLoop anchors (left) and between cKO dOCRs and cKO dLoop anchors (right). ATAC-seq tracks in cKO and floxed neurons are colored in green and purple, respectively, dOCRs are shown as colored bars, dLoops are presented as arcs and the dLoop anchors are highlighted in grey.

Profiling accessible chromatin in cKO vs floxed neurons by ATAC-seq revealed a strong effect of SATB2 on open chromatin landscape (Fig. 1E). Over 28 % of all identified open chromatin regions (OCRs) were differentially accessible between the two genotypes (dOCRs). Of those, 12280/89917 peaks (13.6 %) had greater accessibility in floxed than cKO neurons (Flx dOCRs), whereas 13371/89917 peaks (14.9 %) had greater accessibility in cKO than floxed neurons (cKO dOCRs). A total of 64266 peaks were invariant in both genotypes. Invariant as well as dOCRs were enriched for SATB2 binding sites (invariant OCRs, z-score = 328.14, P < 0.0001; dOCRs, z-score = 19.98, P < 0.0001). While invariant OCRs were enriched for promoter and enhancer chromatin states to similar levels, dOCRs were 6-fold more enriched for enhancers compared to promoters (Fig. 1C). Genomic distribution analysis also showed higher fraction of non-promoter regions (intergenic + intronic + exonic) overlapping with dOCRs compared to invariant OCRs (Fig. 1F). Functional annotation of the genes near invariant OCR revealed that they were predominantly involved in housekeeping functions such as ribonucleoprotein complex biogenesis, ncRNA processing or protein catabolic process (Fig. 1G). By contrast, genes proximal to SATB2-dependent dOCRs were enriched in specialized synapse-related pathways including vesicle-mediated transport in synapse and regulation of membrane potential (Fig. 1G). Therefore, both SATB2 peaks and SATB2-dependent dOCRs are linked to genes encoding neurotransmitter receptors, ion channels, and proteins related to synaptic organization.

Since SATB2 homomeric complexes are predicted to connect distally located DNA binding sites^19^, we next examined the effect of SATB2 deletion on chromatin looping. Loop calling at 5 kb resolution in merged replicate Hi-C contact maps identified 11847 loops (36 % of all loops called in the floxed Hi-C matrix) that were stronger in floxed but weakened/absent in the cKO sample (Flx dLoops). By contrast, 9230 loops (31 % of all loops called in the cKO Hi-C matrix) were detected in cKO but weakened/absent in floxed Hi-C maps (cKO dLoops). A total of 36669 SATB2-independent invariant loops did not differ between the two genotypes. Notably, the anchors of both Flx and cKO dLoops were enriched for SATB2 binding sites (Flx dLoops, z-score = 19.9, P < 0.0001; cKO dLoops, z-score = 18.5, P < 0.0001). Since SATB2 binds predominantly to promoters (Fig. 1A), we stratified dLoops in promoter-based and non-promoter-based by intersecting their anchors with gene promoters. Promoter-based dLoops represented 13.4 % of all chromatin loops in Flx neurons and 11.5 % in cKO neurons. Non-promoter-based dLoops, in which none of the anchors overlapped with a promoter, comprised 22.4 % of chromatin loops in Flx and 19.6 % in cKO neurons. To test for a relationship between dOCRs and dLoops, we further intersected the anchors of dLoops with differential ATAC-seq peaks. This analysis revealed an unexpected correlation between the effects of SATB2 on chromatin accessibility and chromatin looping. The anchors of Flx dLoops were highly enriched for Flx dOCRs, and vice versa, the anchors of cKO dLoops significantly overlapped with cKO dOCRs and were depleted for Flx dOCRs (Fig. 1H-I). This result indicates that the two types of alterations in neuronal 3D epigenome, caused by SATB2 deletion, are tightly coupled.

### SATB2 links regulatory elements to promoters of activity-regulated and cognition-associated genes

Previously described strong effects of SATB2 on gene transcription together with its preferential binding to promoters support a function of SATB2 as a transcription factor (TF), acting directly at promoters^16,30–33^. To examine how SATB2-dependent changes in 3D-epigenome are associated with promoter activity, we first analyzed promoter-centered alterations in chromatin accessibility and looping. The integrative analysis of differential ATAC-seq and Hi-C interaction profiles upon SATB2 deletion revealed two types of SATB2-dependent interactions between distal elements and promoters: dOCR-promoter interactions, in which a dOCR is linked to a gene promoter by a genotype-independent invariant loop (Fig. 2A, Fig. S3A-B), and dLoop-promoter interactions, in which a distal element is connected to a promoter via SATB2-dependent differential loop (Fig. 2B, Fig. S3C-D). To identify dOCR-promoter interactions, we overlapped the anchors of invariant loops, called in both genotypes, with dOCRs and gene promoters. Thereby, 472 genes were assigned to Flx dOCRs and 646 genes to cKO dOCRs. SATB2-driven open chromatin changes and gene expression changes were tightly correlated, since both Flx dOCR- and cKO dOCR-gene sets were enriched for differentially expressed genes (DEGs), previously reported in cKO vs floxed cortical neurons^16^ (Flx dOCR-gene set, OR = 2.3, P = 1.2×10e-14; cKO dOCR-gene set, OR = 2.04, P = 4.1×10e-13). Furthermore, DEGs assigned to Flx dOCRs were expressed at higher levels in floxed neurons, while DEGs assigned to cKO dOCRs were more strongly expressed in cKO neurons (Fig. 2C, Fig. S3A-B). Notably, neuronal activity-regulated genes^34^, which are a subset of DEGs between cKO vs floxed neurons^16^, were strongly overrepresented in the Flx dOCR-but not the cKO dOCR-gene set (OR = 5.56, P < 10e-7), which is consistent with the impaired IEG response observed in cKO neurons^16^. Moreover, we found a strong enrichment for AP-1 TF motifs at Flx dOCRs interacting with promoters. Since AP-1 proteins are key regulators of neuronal activity-driven gene transcription^35^, these findings suggest that the deficient IEG response in cKO neurons is linked to decreased accessibility of AP-1 sites in the regulatory elements of activity-regulated genes. In addition, motifs for MEF TFs, known to be important for synapse formation and elimination^36^, were enriched at promoter-interacting Flx dOCRs. Consistent with this result, we observed an overrepresentation of MEF2C-regulated genes in the Flx dOCR-gene set (OR = 3.3, P < 10e-12)^37^. Salient examples of MEF2C target genes that are also members of Flx dOCR-gene set include *Egr3, Npas4, and Kcna*4^38^. In line with these findings, GO analysis of the Flx dOCRs-gene set revealed strong enrichment for synapse- and memory-linked genes (Fig. 2D). By contrast, GO analysis of the cKO dOCR-gene set identified enrichment of genes associated with forebrain development, regulation of neurogenesis, neuron migration and differentiation (Fig. 2D). Genes known to be highly expressed in neural progenitors but not in mature neurons *such as e*.*g*., *Nrp2, Gdpd5*, and *Gja1* were included in the cKO dOCR-gene set^39–41^. Since these genes are upregulated upon SATB2 loss^16^, SATB2 might be involved in their developmental down-regulation by decreasing chromatin accessibility of the corresponding regulatory elements during late cortical development.

**Figure 2:**
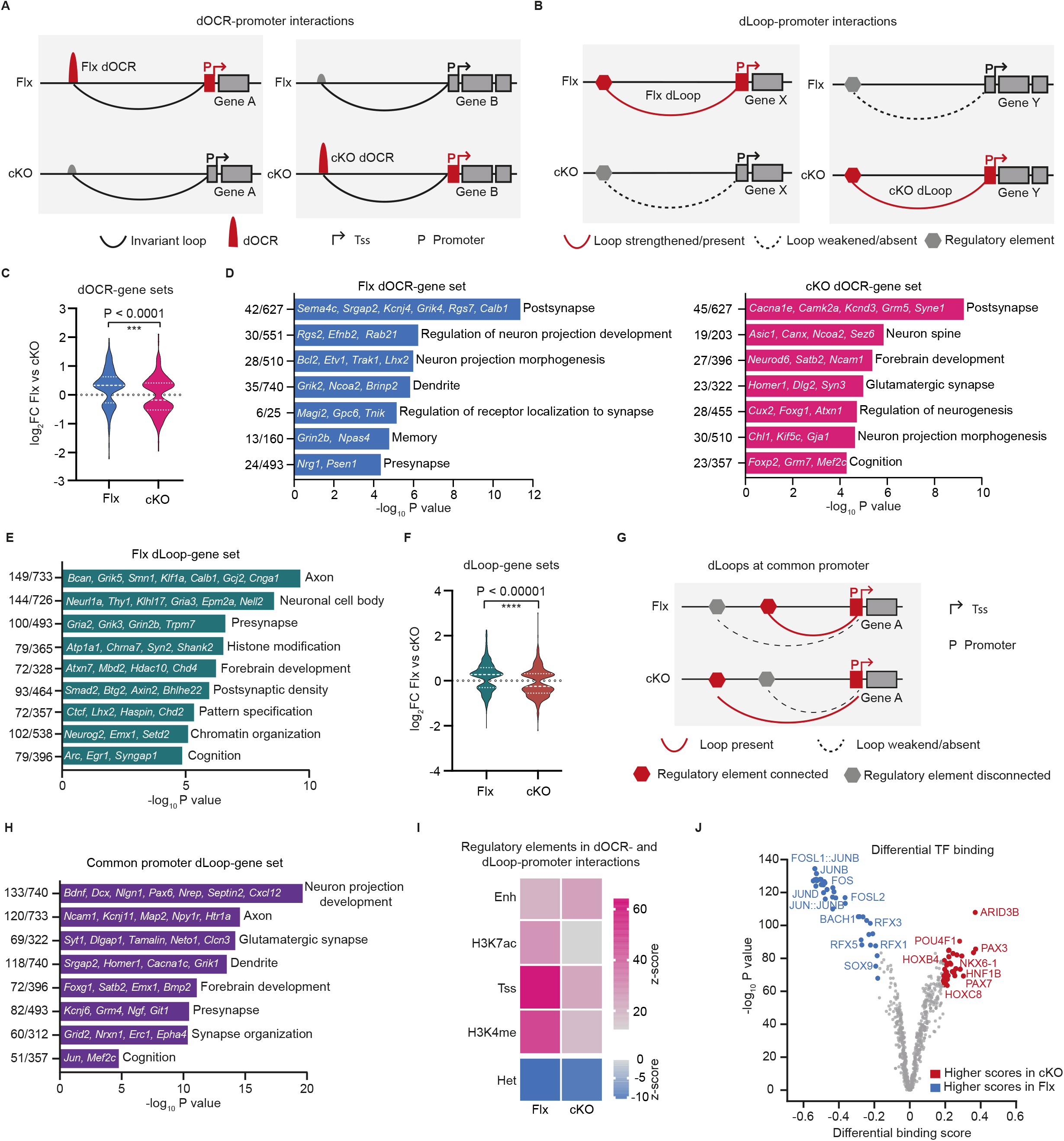
SATB2-dependent regulatory element-promoter interactions involve synapse- and cognition-associated genes. A. Schematic representation of dOCR-promoter interactions: Flx dOCR-promoter interaction (left), cKO dOCR-promoter interaction (right). B. Schematic representation of dLoop-promoter interactions: Flx dLoop-promoter interaction (left), cKO dLoop-promoter interaction (right). C. Fold change distributions of the DEGs assigned to Flx and cKO dOCR-gene sets, respectively. DEGs mapped to Flx dOCR-gene set were more highly expressed in floxed neurons and vice versa, DEGs mapped to cKO dOCR-gene set were more highly expressed in cKO neurons. P value is calculated using non-parametric Mann-Whitney test. D. GO analysis of Flx dOCR-gene set (left panel) and cKO dOCR-gene set (right panel). E. GO analysis of Flx dLoop-gene set. F. Fold change distributions of the DEGs assigned to Flx and cKO dLoop-gene sets, respectively. P value is calculated using non-parametric Mann-Whitney test. G. Schematic representation of dLoop-promoter interactions at common promoter. H. GO analysis of common promoter dLoop-gene set. I. Enrichment analysis of distal using ChromHMM-infered epigenomic states and histone marks (P0 forebrain). Color bar shows the z-score value, as calculated by permutation tests with regioneR, *P < 0.001. J. Differentially bound TFs across ATAC-seq peaks at promoter-interacting distal elements, identified by TOBIAS. TF motifs with higher footprinting score in Flx and cKO samples are shown in blue and red, respectively.

To identify dLoop-promoter interactions, we intersected the anchors of Mustache-called dLoops with promoters (Fig. 2B, Fig. S3C). Thereby, 2978 and 2133 genes were assigned to Flx dLoops and cKO dLoops, respectively. To gain insight into their biological functions, we first examined the associated GO terms. Notably, the Flx dLoop-gene set was specifically and highly enriched for axonal and synaptic genes, as well as for genes linked to cognition and forebrain development (Fig. 2E). By contrast, the cKO dLoop-gene set did not significantly associate with any GO term and thus appeared to be mostly random. Noteworthy, although the Flx dLoop- and Flx dOCR-gene sets were defined by different underlying molecular mechanisms and differed in their composition, we noticed striking functional similarities between them. Like the Flx dOCR-gene set, also the Flx dLoop-gene set overlapped significantly with SATB2-dependent DEGs^16^ (OR = 1.76, P < 10e-16), neuronal activity-regulated genes^34^ (OR = 1.96, P < 10e-4) and MEF2C-regulated genes^37^ (OR = 1.94, P < 10e-12). Furthermore, in both cases the DEGs within the gene sets were expressed at higher levels in floxed neurons (Fig. 2D-F, Fig. S3C), and the ATAC-seq peaks within promoter-interacting distal targets of Flx dOCR-genes and Flx dLoop-genes were enriched for AP1 and MEF TF motifs. Intriguingly, in 43 % of Flx and 52 % of cKO dLoop-promoter interactions the same promoter was connected to alternative targets in cKO vs floxed neurons (Fig. 2G, Fig. S3D). This alternative usage of regulatory elements resulted in differential gene expression since genes within this gene set overlapped significantly with the DEGs in cKO vs Flx cortical neurons^16^ (OR = 2.24, P < 10e-16). GO analysis demonstrated again a strong enrichment of neuron-specific and synaptic terms in this common promoter dLoop-gene set (Fig. 2H). Prominent examples of genes within this set include *Rgs7, Cacna1c*, and *Srgap2*.

To characterize the epigenetic features of the distal elements linked to promoters by dOCR- and dLoop-promoter interactions, we leveraged ChromHMM defined chromatin states, neonatal forebrain H3K27Ac and H3K4me3 ChIP-seq data^27,42,43^, and the SATB2 binding sites, defined in this study. As expected, targets interacting with promoters in a SATB2-dependent manner were enriched for SATB2 peaks (Flx, z-score = 37.77, P < 10e-6, cKO, z-score = 30.28, P < 10e-6). Since enhancer and promoter states were also strongly enriched (Fig. 2I), we conclude that dOCR- and dLoop-promoter interactions are mostly of enhancer-promoter or promoter–promoter type, potentially forming neural enhancer-promoter aggregates^44^. Notably, there were significant differences in TF binding between Flx vs cKO targets^45^. Motifs for AP-1 (e.g. FOS, JUNB, JUND, JUN and FOSL1/2) and RFX (RFX1-5) families of TFs had the highest footprinting score at Flx distal elements, whereas motifs for homeobox TFs (e.g. HNF1B, PAX7, PAX3, POU4F1) and ARID3B showed elevated footprinting at cKO distal elements (Fig. 2J).

In summary, the integration of Hi-C and ATAC-seq data revealed that in cortical neurons SATB2 organizes distal element-promoter interactions by at least two different mechanisms: 1) differential accessibility of enhancers within invariant chromatin loop interactions, and 2) differential chromatin looping between promoters and distal enhancers. Both types of SATB2-dependent 3D epigenome changes tightly correlated with gene expression and affected with remarkable specificity genes with specialized synaptic functions.

### SATB2 shapes the 3D genome architecture of cortical neurons at multiple hierarchical levels

Having demonstrated a role of SATB2 in mediating distal element-promoter loops, we next asked if SATB2 modulates 3D chromatin structure at larger genomic scales. To gain insights into the global genome folding pattern upon SATB2 loss, we first explored distance-dependent chromatin interaction frequencies. Genome-wide average contacts decayed stronger with distance in cKO vs floxed Hi-C contact matrices at all distances (Fig. 3A), indicating a lower level of chromatin compaction in cKO compared to floxed neurons^46^. Consistent with this finding, we detected a weakened compartmentalization in cKO neurons, mostly within the inactive (B), but also within the active (A) compartment, compared to floxed samples (Fig. 3B). Collectively, these genome-wide aggregate analyses revealed global changes in 3D genome architecture upon SATB2 deletion that are comparable in effect-size to those described after loss-of-function of well-established spatial genome organizers, such as CTCF and cohesin^47^.

**Figure 3:**
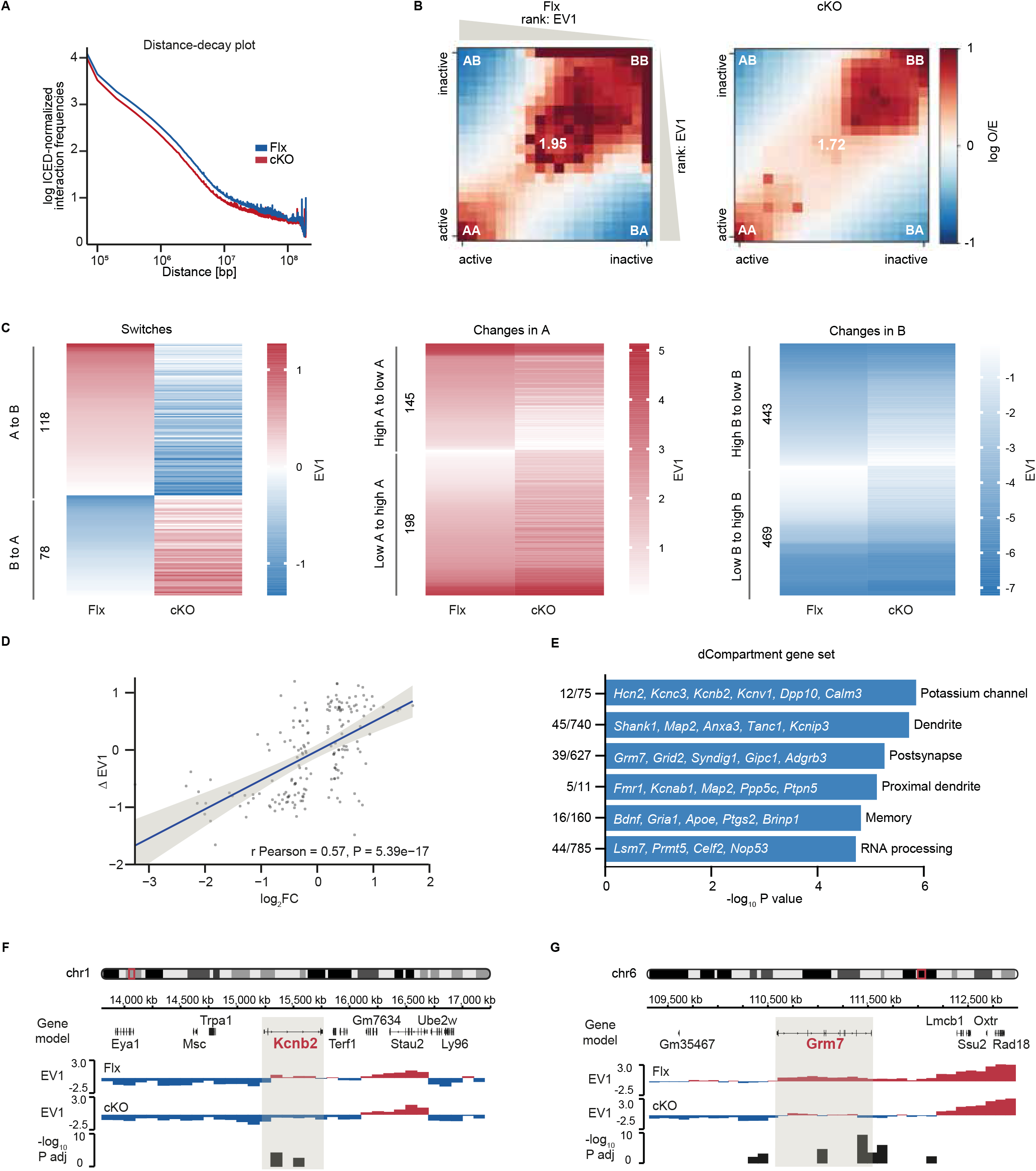
SATB2 deletion causes global reduction in both intra-chromosomal contact frequency and long-range compartmentalization strength. A. Chromosome-averaged distance-decay plots illustrating intra-chromosomal contact frequency as a function of genomic distance in cKO and floxed Hi-C matrices. B. Saddle plots showing compartmentalization of the genome in cKO and Flx samples. Shown are the average observed/expected (O/E) values of contact enrichment between pairs of 100 kb bins arranged by their compartment signal strength (EV1) from highest (most A-like) to lowest (most B-like). Numbers in white represent the compartment strength. C. Heatmaps illustrating changes in compartment state upon SATB2 deletion. Left, compartment switches (from inactive B-to permissive A-compartment and vice versa), right, shifts in compartment score within the same compartment. D. Correlation between changes in compartment score (ΔEV1) and expression (log2FC) for DEGs located within differential compartments. E. GO analysis of genes mapped to SATB2-dependent differential compartments (dCompartment gene set). F. Representative IGV browser tracks showing compartment switch (left) and compartment strength change (right) in cKO vs floxed neurons. Shown are EV1 values and –log_10_ P adj values of the differential compartment calls (100 kb resolution).

We next identified specific genomic regions that undergo compartment changes. Differential compartment analysis^48^ at 100 kb resolution revealed a total of 1451 pairwise significant differential compartments between cKO and floxed neurons (FDR < 0.01), covering 5.6 % of the genome. Compartment changes were grouped into switches (196 significant changes from A to B and from B to A compartment) and strength changes (912 in B compartment strength and 343 in A compartment strength) (Fig. 3C). For the 631 genes mapped to differential compartments (dCompartment gene set), shifts in the compartment score (ΔEV1) were positively correlated with changes in gene expression (Fig. 3D). The dCompartment gene set was highly enriched in postsynapse-, dendrite- and memory-related GO terms, again demonstrating the high functional specificity of SATB2-dependent effects on chromatin structure in pyramidal neurons, this time at compartment scale. Examples of synaptic plasticity-linked genes undergoing compartment shift/strength change include *Bdnf, Grm7, Fmr1, and Kcnb2* (Fig. 3F). Notably, several SZ risk prioritized genes^49^, including *Csmd1, Lrrc4b, Ptprd, and Epn2* (the later also associated with intelligence^50^) were also among the genes mapped to differential compartments.

Next, we tested for rearrangements in TADs upon SATB2 deletion, using a method for TAD calling based on spectral clustering^51^. The results showed similar number and average width of TADs and subTADs in floxed vs cKO Hi-C matrices (Fig. S4A-B). However, the differential analysis of TAD boundaries^52^ revealed changes in 1203 boundaries between cKO and floxed Hi-C datasets, representing 12.7 % of all detected TAD boundaries. Different types of TAD boundary changes were identified, including complex, split, merge, shifted, and strength changes (Fig. 4A-B). The genes located within 100 kb regions flanking differential boundaries overlapped significantly with DEGs between cKO vs floxed cortical neurons (OR = 1.55, P = 0.00002). Furthermore, differential TAD boundaries were associated with SATB2-dependent gene regulation since genes nearby Flx TAD boundaries (Flx dBoundary gene set) were expressed at higher levels in floxed neurons, while genes close to cKO TAD boundaries were expressed at higher levels in cKO neurons (Fig. 4C). The Flx dBoundary gene set (504 genes) was strongly enriched for GO terms related to synaptic transmission and neuron projection development (Fig. 4D). By contrast, the genes nearest to cKO TAD boundaries (310 genes) did not show significant enrichment for any GO term. Notably, Flx and cKO specific TAD boundaries further differed in their epigenetic profile, defined by ChromHMM-predicted chromatin states^42^. While Flx TAD boundaries were enriched for genomic annotations known to be associated with TAD boundaries, e.g., promoters and CTCF binding sites^53–55^, cKO TAD boundaries did not show enrichment for any of these features (Fig. 4E). Previous data have demonstrated that cell type-specific TAD boundaries are likely to harbor genes that are functionally relevant to the corresponding cell type^56^. Considering the epigenetic features of Flx TAD boundaries and the enrichment of synapse-related terms in the genes nearby, our results suggest that SATB2 determines pyramidal neuron-specific TAD borders with impact on expression of neighboring genes, exerting specific neuronal functions. Next, we analyzed the characteristics of non-promoter-based dLoops (nonP-dLoops) that represent more than half of SATB2-dependent differential loops and with a median length of ∼355 kb belong to the sub-megabase level of genome organization^57^. Unexpectedly, we found an enrichment of CTCF peaks at anchors of nonP-dLoops (Flx nonP-dLoops, z-score = 60.41, P < 10e-6, cKO nonP-dLoops, z-score = 62.06, P < 10e-6) (Fig. 4F). CTCF has been established as a major architect of 3D genome topology at the sub-megabase level^58^, therefore we investigated potential functional links between CTCF and SATB2. Since our previous interactome analyses in cortical neurons^12^ did not provide evidence for physical CTCF-SATB2 interactions, we tested the potential of SATB2 and CTCF to cooperate indirectly, via common co-interactors. A significant overlap was detected between SATB2 interactors in cortical neurons^12^, CTCF interactors^61,62^ and CTCF loop participants/co-localizing factors^60^ (Fig. 4G). Given that CTCF-dependent loops appear to be regulated by multi-component CTCF complexes^60,63^, the effects of SATB2 on nonP-dLoops are likely to be mediated via modulation of the strength of CTCF loops in complexes containing both proteins.

**Figure 4:**
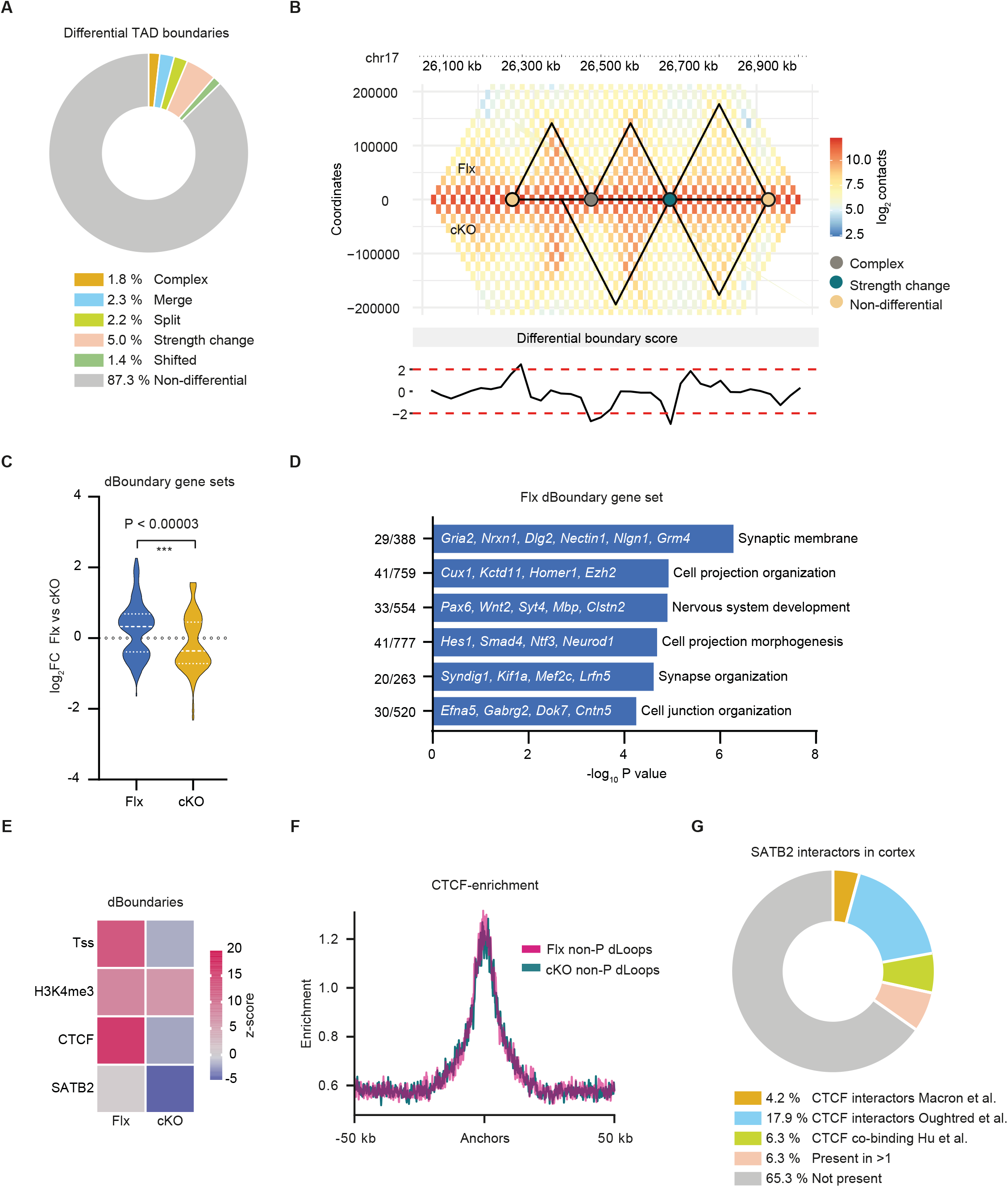
TAD boundary changes and non-promoter-based differential loops occur at genomic regions occupied by CTCF. A. Pie chart showing the proportion of different types of TAD boundary changes between cKO and floxed samples. B. Representative example of differential boundaries between cKO and floxed neurons (chr17:26033500–26983500). Outlined TADs were identified by using SpectralTAD TAD caller at 25 kb resolution. C. Fold change distributions of DEGs near differential TAD boundaries. *P* value is calculated using non-parametric Mann-Whitney test. D. GO analysis for genes nearby Flx-specific TAD boundaries (Flx dBoundary gene set) as defined by GREAT^59^ (association rule: basal + extension: 5 kb upstream, 1 kb downstream, and 100 kb max extension). E. Enrichment analysis of differential TAD boundaries over SATB2 binding sites, CTCF binding sites, H3K4me3 peaks, and promoter states, inferred by ChromHMM (P0 forebrain). Color bar shows the z-score, as calculated by permutation tests with regioneR, for all enrichments, P < 0.001. F. Average CTCF ChIP signal (ENCODE ChIP-seq from P0 forebrain) over anchors of non-P-based differential loops (nonP-dLoops). G. SATB2 protein interactors overlap significantly with a set of CTCF co-binding factors^60^ / CTCF protein interactors^61,62^. Fisher’s exact test was used for statistical analysis, P < 10e-16, OR = 16.04.

In conclusion, our data establish SATB2 as a cell type-specific determinant of pyramidal neuron 3D chromatin structure, acting not only at chromatin loop level, but also at multiple larger scale hierarchical levels. These findings demonstrate a previously unexpected role of SATB2 as an architectural protein, influencing long-range chromatin interactions.

### SATB2-dependent FIREs and super-FIREs harbor cognition-associated genes

Frequently Interacting Regions (FIREs) are tissue- and cell type-specific local interaction hotspots that are enriched for active *cis*-regulatory elements^64,65^. Given the highly restricted, pyramidal neuron-specific expression pattern of SATB2, we tested if SATB2 acts as a FIRE determinant in cortical neurons. A total of 6669 and 5428 FIREs were detected in floxed and cKO Hi-C contact matrices, respectively. To identify SATB2-dependent FIREs, we used a statistical cutoff based on the differences in FIRE scores between cKO and floxed neurons^64^, thereby detecting 99 Flx- and 15 cKO-specific FIREs (Fig. 5A). Consistent with previous findings^65^, Flx FIREs were enriched for enhancer states (z-score = 4.39, P < 0.0001). Furthermore, Flx FIREs were specifically enriched for Flx dOCRs, indicating a correlation between SATB2 effects on FIREs and chromatin accessibility (z-score = 3.67, P < 0.002). By contrast, cKO FIREs did not show any enrichment for active chromatin. Contiguous FIREs are known to form clusters, called super-FIREs, which have the most significant local interaction frequency^65,66^. By filtering out super-FIREs common to cKO and Flx neurons on a genome-coordinate level, we identified 180 Flx- and 93 cKO-specific super-FIREs. To investigate how FIREs and super-FIREs are associated with SATB2-driven gene expression, we intersected them with genic annotations, thereby assigning 49 and 123 genes to Flx FIREs and superFIREs, and 8 and 89 genes to cKO FIREs and super-FIREs, respectively. Genes mapped to Flx FIREs/super-FIREs showed higher expression levels in Flx compared to cKO neurons and were strongly enriched for DEGs (FIREs, OR = 5.1, P < 10e-7, super-FIREs, OR = 3.62, P < 10e-10) (Fig. 5B). In fact, all DEGs mapped to Flx FIREs and 48/49 DEGs mapped to Flx super-FIREs were upregulated in floxed compared to cKO neurons. GO analysis of Flx FIRE/super-FIRE gene sets showed significant enrichment of synapse-related terms as well as terms associated with behavior, learning or memory, and cognition (Fig. 5C). By contrast, the 8 genes located in cKO FIREs and 89 genes in cKO super-FIREs did not show significant overrepresentation of any GO category.

**Figure 5:**
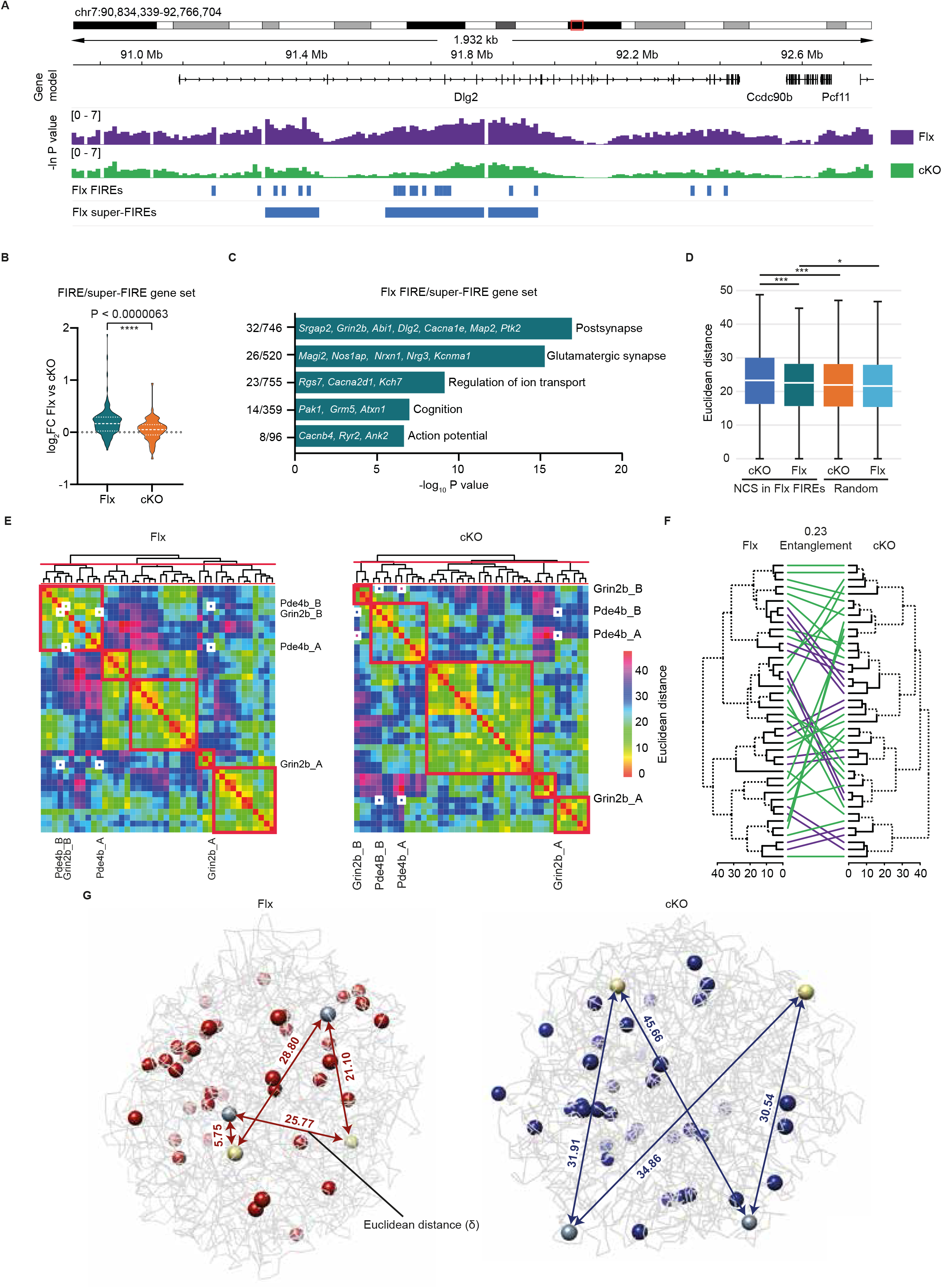
SATB2-dependent FIREs and super-FIREs overlap with cognition-related genes. A. Representative IGV browser tracks illustrating an example of Flx FIREs and super-FIREs. The significance of FIRE scores, as calculated by one-sided Z-test, in floxed and cKO neurons is depicted in green and purple, respectively. Flx FIREs and super-FIREs are presented as blue bars under the significance tracks. B. Genes located in Flx FIREs/super-FIREs were more highly expressed in floxed compared to cKO neurons. P value is calculated using non-parametric Mann-Whitney test. C. GO analysis of genes mapped to Flx FIREs/super-FIREs. D. Box plots showing the pairwise Euclidean distances (ED) between 43 domains harboring Flx FIREs/super-FIRE genes with human orthologues mapped to NCS meta-loci in Flx vs cKO 3D genome models (P = 3.27e-13, Wilcoxon signed rank test). Shown are the pairwise ED between randomly permuted sets of domains of the same size (n = 1000 permutations) in Flx vs cKO 3D genome models (P =3.06e-09 NCS in Flx FIREs vs random domains, cKO 3D model; P = 0.016, NCS in Flx FIREs vs random domains, Flx 3D model, Wilcoxon rank-sum tests). E. Heatmaps of pairwise ED between the 43 domains harboring Flx FIREs/super-FIRE genes with human orthologues mapped to NCS meta-loci in Flx vs cKO 3D genome models. Red squares depict five Euclidean hot spots (clusters of domains that are spatially close together) in Flx and cKO sample. White squares show domains harboring *Grin2b* and *Pde4b* alleles, a representative pair of Flx FIREs/super-FIREs genes having human orthologues mapped to NCS meta-loci. F. Comparison between Flx and cKO dendograms of the 43 domains harboring Flx FIREs/super-FIRE genes with human orthologues mapped to NCS meta-loci. “Unique” nodes, with a combination of labels/items not present in the other tree, are highlighted with dashed lines. Green colored lines connect domains that belong to different subtrees in the two dendogrames. G. *In silico* Chrom3D models of cortical neuron spatial genome, Flx 3D genome model (left), cKO 3D genome model (right). Colored beads correspond to the 43 domains harboring Flx FIREs/super-FIRE genes with human orthologues mapped to NCS meta-loci, dark khaki and slate grey beads represent domains, harboring *Grin2b* and *Pde4b* alleles, respectively. Pairwise ED are reported as numbers above the lines connecting dark khaki and slate grey beads.

Considering the strong enrichment of cognition-related GO terms in the Flx FIRE/super-FIRE gene sets, we next tested if they include mouse orthologues of the human genes, mapped to General Cognitive Ability (GCA) and Non-Cognitive Skills (NCS) meta-loci^67^. A meta-locus is defined as a set of linkage disequilibrium-independent genomic regions that show similar local genetic correlation profiles across several psychopathology traits and GCA and NCG, respectively^67^. In total, 13/45 Flx FIRE genes and 29/114 super-FIRE genes were identified as the mouse orthologues of GSA “driver” genes (ORs = 9.41 and 7.74, P < 10e-8 and 10e-14). Likewise, 12/45 Flx FIRE genes and 14/114 super-FIRE genes were identified as the mouse orthologues of NCS “driver” genes (ORs = 16.15 and 5.76, P < 10e-10 and 10e-16). Examples of Flx FIRE/super-FIRE-mapped genes with human orthologues harbored in GCA meta-loci include *Anks1b, Gphn, Rere*, and *Plcb1*, whereas *Dlg2, Grin2b, and Nrxn1* are located in NCS meta-loci. A substantial fraction of Flx FIRE/super-FIRE genes with human orthologue in GCA/NCS meta-loci were also associated with autism spectrum disorder (ASD)^68^ (GCA, 13/37; NCS, 11/21), developmental brain disorders^69^ (GCA, 10/37; NCS, 8/21), and intellectual disability (ID)^70^ (GCA, 10/37; NCS, 5/21).

Gene sets harbored in NCS and GCA meta-loci are coordinately regulated over lifespan and share specific biological pathways^67^. Proximity in intra-nuclear positioning and/or increased connectivity between regulatory elements have been suggested as contributing mechanisms for gene co-regulation^71^. Given the extensive overrepresentation of NCS and GCA driver genes in Flx-FIREs/super-FIRE gene sets, we reasoned that SATB2 might regulate the spatial distance between them within the virtual 3D sphere of the cortical neuronal nucleus. To test this hypothesis, we modelled chromatin conformation in 3D and visualized nuclear topography by Chrom3D-based Monte Carlo simulations^72^. First, we created 3D models by using a consensus set of Chrom3D TAD domains, called at 50 kb resolution, and Hi-C data from floxed vs cKO neurons. These models consisted of 3478 chromatin bead domains called for the diploid genome, averaging 1.42 Mb in length. Next, we mapped Flx FIRE/super-FIRE-associated genes with human orthologues in GCA and NCS meta-loci to Chrom3D domains, resulting in 43 and 69 domains, respectively. Computing pairwise Euclidean distances between these domains resulted in significantly higher proximity and shorter Euclidean distances in floxed compared to cKO 3D models (Fig. 5D-G, Fig. S4C). Extending this analysis to all NCS and GCA driver genes i.e., beyond the subsets mapped to Flx FIREs/super-FIREs, we obtained similar results of closer proximity and shorter Euclidean distances in floxed versus cKO 3D models (P = 2.2e-16 for domains mapped to NCS and GCA, Wilcoxon signed-rank tests). In addition, we performed hierarchical clustering of the domains harboring Flx-FIRE/super-FIRE genes with NCS/GCA driver gene orthologues, based on pairwise Euclidean distances derived from floxed vs cKO 3D models. Cutting the trees at 0.90 quantile resulted in clusters of domains that are confined within close proximity within the 3D nuclear space (Euclidean “hot spots”)^73^ (Fig. 5E, Fig. S4D). Notably, although Flx and cKO dendograms consisted of the same total number of Euclidean “hot spots”, there was a substantial re-shuffling of domains from cKO into floxed Euclidean “hot spots” and vice versa (Fig. 5E-F, Fig. S4D-E). The dissimilarity between cKO and Flx dendograms was further supported by low values of Cophenetic and Baker correlation coefficients, revealing low level of similarity between the two trees. The weak correlation between dendogrames was not restricted to the subsets of NCS and GCA driver genes mapped to Flx-FIREs/superFIREs, but was also observed when the complete NCS and GCA driver gene sets were compared. Therefore, SATB2 affects distances between NCS and GCA driver genes both inside and outside of local interaction hotspots (FIREs/super-FIREs) in pyramidal neurons. These data establish SATB2 as a critical regulator of the nuclear topography of NCS/GCA driver genes, related to both cognitive dimensions and psychopathology.

### SATB2-dependent 3D epigenome changes contribute to cognitive ability and risk for neurodevelopmental and neuropsychiatric disorders

To explore the functional relevance of the genes affected by SATB2-dependent 3D epigenome changes, we pooled all sets of genes with SATB2-dependent 3D chromatin structure into a single gene set (SATB2 3D genome set). The following gene sets were combined: dCompartment gene set, Flx dBoundary gene set, Flx and cKO dOCR-gene sets, Flx dLoop-gene set, common promoter dLoop-gene set, and Flx FIRE/super-FIRE gene set. Given the lack of significant GO term enrichment in the cKO dBoundary gene set, cKO dLoop-gene set, and cKO FIRE/super-FIRE gene set, these were considered as representing randomly gained effects upon SATB2 loss and thus excluded from the analysis. We next tested if SATB2 3D genome set, as a whole, overlaps with genes comprising relevant neuron- and cognition-specific GO categories (Fig. 6A). Between 46 % and 50 % of all genes belonging to these categories were found to be linked to SATB2-dependent 3D genome architecture, resulting in a highly significant overrepresentation (Fig. 6A). By contrast, no overrepresentation of the SATB2 3D genome gene set was detected for control GO categories and a significant depletion was observed in some of them (Fig. 6A). Remarkably, the individual gene sets, defined by SATB2-dependent effects at individual 3D genome hierarchical levels, showed little overlap between each other. More than 70 % of the genes within the SATB2 3D genome set were exclusively represented in only one hierarchical level-specific gene set (Fig. 6A). Therefore, SATB2 selectively influences the 3D chromatin structure of a large proportion of genes with defined cognition-related function by operating at various chromatin architectural levels and affecting essentially different subsets of genes on each level.

**Figure 6:**
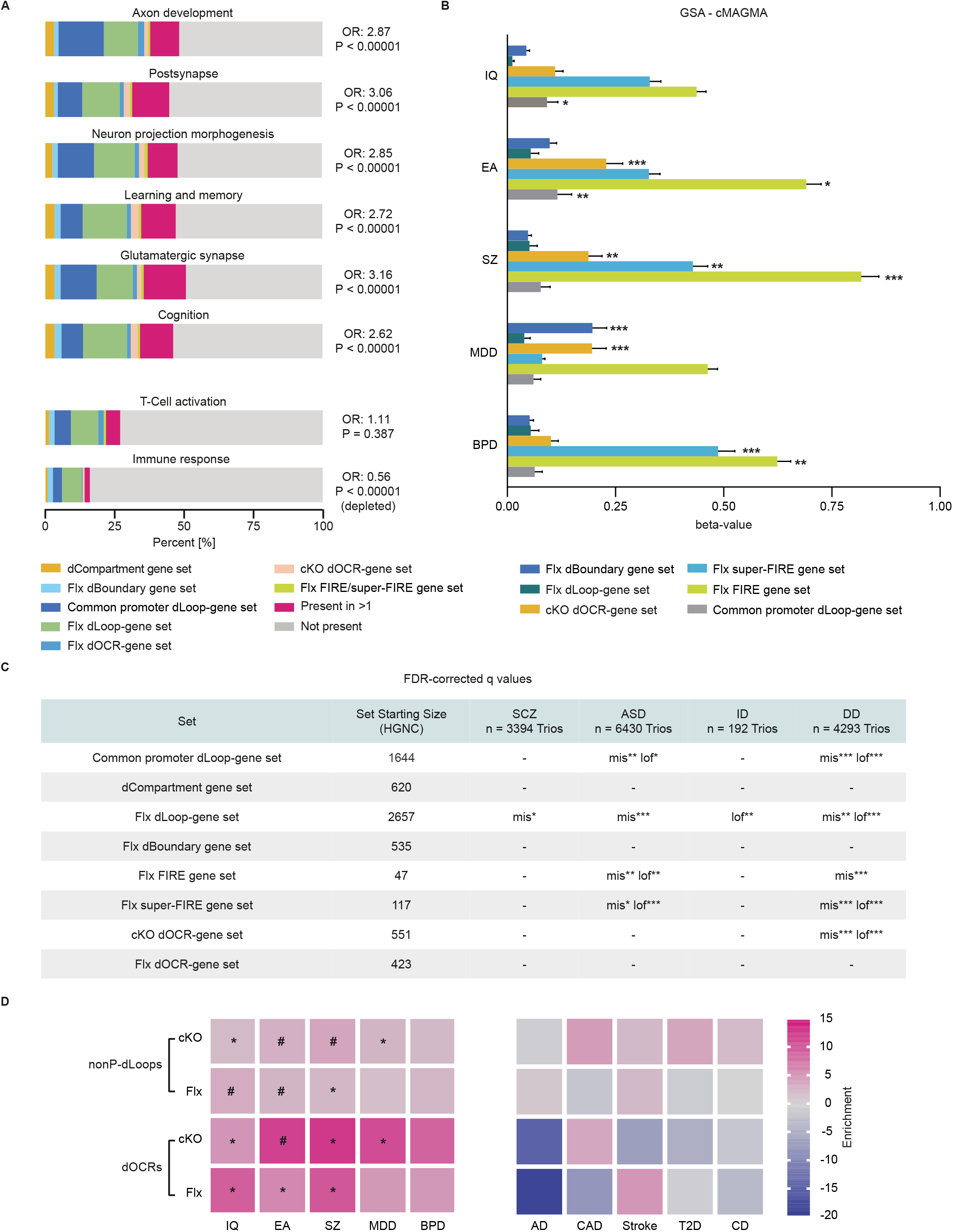
Genes and non-coding regions linked to SATB2-dependent 3D genome changes contribute to cognitive ability and risk for neuropsychiatric disorders. A. Neuron- and cognition-specific GO categories are enriched for genes affected by SATB2 3D epigenome remodeling. Bar charts showing the proportion of these GO categories occupied by the gene sets representing the effects of SATB2 at individual 3D genome hierarchical levels. P values were calculated using Fisher’s exact test, significance level after Bonferoni correction of multiple testing = 0.005, OR, odds ratio. B. MAGMA-GSA of SATB2 3D genome sets in EA, IQ, SZ, BP, and MDD. Beta values (effect sizes) are plotted on the x-axis with P values shown beside each bar. *P < 0.05, **P < 0.01, ***P < 0.001. Horizontal bars indicate standard error. C. Analysis of SATB2 3D genome sets using data on *de novo* mutations, identified in SZ, ASD, ID, and DD, mis = missense mutations; lof = loss-of-function mutations; * P _Bonferroni_ < 0.05; **P _Bonferroni_ < 0.01; ***P _Bonferroni_ < 0.001. Shown are total number of genes (per set) with a human orthologue (HGNC, HUGO Gene Nomenclature Committee). D. Enrichment of common risk variants for brain-related and non-psychiatric/non-brain related GWAS traits in the human orthologues of dOCRs and nonP-dLoop anchors. Enrichment values are plotted as heatmap. #, P values significant for enrichment in LDSR after Bonferoni correction of multiple testing across all tests, *, P values nominally significant for enrichment.

We next examined if genes within the individual SATB2 3D gene sets contribute to human psychiatric disorders and cognitive phenotypes. We first tested if common variation in the corresponding human orthologues is associated with major psychiatric traits by using gene set analysis (GSA). To assign SNPs to cognate genes, we implemented MAGMA^74^ and H-MAGMA^75^, the latter leveraging chromatin interaction profiles from human brain tissue for gene mapping. MAGMA-GSA showed an enrichment for genes associated with educational attainment (EA) in the following gene sets: common promoter dLoop-gene set, cKO dOCR-gene set, and Flx FIRE/super-FIRE gene sets (Fig. 6B). Enrichment for genes associated with cognitive ability/human intelligence (IQ) was detected in the common promoter dLoop-gene set. Enrichment for genes associated with SZ and bipolar disorder (BPD) was found in Flx FIRE/super-FIRE gene sets and cKO dOCR-gene set. The latter gene set was also enriched for genes associated with major depression disorder (MDD). H-MAGMA-GSA using gene-SNP pairs based on cortical neuronal Hi-C^64^ and fetal brain Hi-C^76^ revealed similar enrichments. To examine if the enrichments detected for EA, IQ, SZ, BPD, and MDD are a property of polygenic phenotypes in general, we included several control phenotypes covering both brain-related and non-brain related diseases, such as Alzheimer’s disease, stroke, cardiovascular disease, Crohn’s disease, and type 2 diabetes. No enrichment was observed for any of the five phenotypes tested. Furthermore, the cKO dBoundary gene set, cKO dLoop-gene set and cKO FIRE/super-FIRE gene set, likely derived from non-physiological, gained effects upon SATB2 loss, also did not show enrichment for genes associated with psychiatric disorders and/or cognitive ability, except the cKO dBoundary gene set, which was marginally enriched for genes associated with SZ.

Next, we examined if the individual SATB2 3D genome subsets are enriched for *de novo* mutations, reported in trio-based studies of SZ^77,78^, ASD^79^, intellectual disability^80^, and developmental disorders (DD)^81,82^. We tested for enrichment of missense (Mis) and loss-of-function (LoF) mutations, while synonymous (Syn) mutations served as a control, as they are unlikely to be pathogenic. As an additional control, we analyzed control trios and trios made from unaffected siblings. We found that the Flx dLoop-gene set, the common promoter dLoop-gene set, and the Flx FIRE/super-FIRE gene set were significantly enriched for LoF and/or Mis mutations reported specifically in ASD and DD patients (Fig. 6C). Furthermore, the cKO dOCR-gene set was enriched for both LoF and Mis mutations found in the DD trios. Analysis of Syn mutations in all four disorders did not reveal significant enrichment at FDR < 0.01 in any of the individual subsets. Likewise, both control trios and trios made from unaffected siblings showed no enrichment for *de novo* mutations in any of the SATB2 3D genome subsets. Genes linked to 3D epigenome alterations gained in cKO were also not enriched for *de novo* mutations reported in any of the four trio-based studies.

Finally, we tested if SATB2-dependent alterations in chromatin accessibility or non-promoter centered looping are informative for non-coding genetic variation linked to neuropsychiatric diseases. To this aim, we identified orthologous regions in the human genome of dOCRs and nonP-dLoops by liftover analysis. Importantly, the human orthologues of mouse dOCRs were enriched for enhancer states identified in human frontal cortex^83^, suggesting a potentially conserved function (cKO dOCR, z-score = 13.85, P < 10e-6; Flx dOCR, z-score = 11.50, P < 10e-6). Next, we performed stratified linkage disequilibrium score regression (sLDSC) analysis to investigate if common genetic variants, associated with cognitive ability or risk for neuropsychiatric disorders, are enriched in the human orthologues dOCRs and nonP-dLoop anchors. The results showed a significant enrichment of SNP-based heritability for EA, IQ, and SZ, out of five psychiatric and cognition-related traits tested, in the human orthologous regions of mouse dOCRs and/or nonP-dLoop anchors (Fig. 6D). By contrast, no enrichment in SNP heritability was observed in control non-psychiatric/non-brain related traits and disorders tested (Fig. 6D). The specificity of the observed enrichments, i.e. only for EA, IQ, and SZ, and not for the other two neuropsychiatric disorders, is consistent with the existing molecular genetic evidence for overlap between general cognitive ability and SZ^67,84^, and points to a link between SATB2-directed 3D epigenome remodeling and the common genetic underpinnings of psychiatric and cognitive phenotypes.

Taken together, our data indicate that SATB2-dependent 3D-epigenome alterations, in particular changes in chromatin accessibility, chromatin loops, and FIREs/super-FIREs, are part of the genetic mechanisms contributing to human cognitive function, neuropsychiatric (SZ and BPD) and neurodevelopmental (ASD and DD) disorders.

## Discussion

Our work identifies 3D genome remodeling by SATB2 as a novel mechanism that balances synaptic activity and gene expression within the highly specialized nucleus of postmitotic pyramidal neurons. With remarkable specificity, SATB2 influences the chromatin structure of 46 to 50 percent of all genes that are functionally associated with neuronal plasticity and/or cognition. These SATB2-dependent effects occur at all known 3D genome architectural levels and closely correlate with the expression of the affected genes.

Activity-regulated gene expression is a major mechanism contributing to long-lasting modifications of neural functions, including learning and memory^35^. Neural IEGs comprise a diverse set of genes with a broad structural and functional repertoire. The exact mechanisms orchestrating the coordinate expression of such a heterogeneous set of genes upon neuronal activation remain to be fully established. Previous studies have highlighted the importance of the interplay between chromatin occupancy and chromatin interaction changes, occurring upon synchronous or sparse activation *in vivo*, or during memory formation and recall^85,86^. Altered occupancy of AP1-binding sites at distal regulatory regions of activity-induced genes has been pinpointed as a part of the epigenomic signature of neuronal activation^86^. Our study establishes a prominent role of SATB2 in this process by demonstrating that SATB2-dependent dOCR- and dLoop-promoter interactions occur at a large fraction of activity-regulated and cognition-associated genes. Furthermore, we find an enrichment for AP1-binding sites at promoter-interacting distal anchors of SATB2-dependent differential loops. Thus, our data suggest that the impaired induction of activity-regulated genes in SATB2-deficient neurons^16^ is likely caused by two different mechanisms: reduced accessibility of AP1 sites at distal regulatory elements or weak/absent chromatin loops connecting AP1-bound distal regulatory elements to the promoters of these genes.

Our analysis identifies extensive and strong effects of SATB2 on chromatin accessibility genome-wide. Approximately one third of all detected OCRs in cortical neurons are SATB2-dependent. We distinguish three distinct types of dOCRs, based on their genomic annotation: proximal to promoters, interacting with promoters via invariant loops, and overlapping with anchors of SATB2-dependent non-promoter-based differential loops. By integrating Hi-C, ATAC-seq and transcriptome data^16^, we uncover a function of the first two dOCR-types in local, promoter-based interactions. For genes linked to these dOCRs, alteration in chromatin accessibility, likely caused by SATB2-recruited histone-modifying complexes^87,88^, closely parallels the bidirectional change in gene expression observed upon SATB2 loss. For example, we observe increased chromatin accessibility at dOCRs linked by invariant loops to genes that are highly expressed in neural progenitors^39–41^ and are likely downregulated in SATB2-expressing neurons during late cortical development. By contrast, activity-regulated genes that are down-regulated in cKO neurons are marked by dOCRs with decreased accessibility in cKO. The third and very abundant type of dOCRs is found at anchors of promoter-independent but SATB2-dependent differential loops. These dOCRs likely serve an architectural role in chromatin organization, consistent with recent observations that chromatin state is primarily responsible for chromatin interactions within compartmental domains^89,90^.

Beyond chromatin looping and chromatin accessibility, we uncover unpredicted broader effects of SATB2 on 3D genome folding^90^. Whereas depletion of CTCF, the best-characterized 3D genome organizer, affects only chromatin looping and TAD insulation without influencing higher-order genomic compartmentalization^91^, SATB2 ablation impacts on multiple hierarchical levels, including compartments, TAD boundaries, FIREs, and also affects global chromatin compaction and compartmentalization. Another important aspect of SATB2 function in 3D genome folding, besides its versatility, is the remarkable level of specificity. At all hierarchical levels, the affected loci are not random but involved in highly specialized neuronal processes, including synaptic signaling, regulation of action potential or learning and memory. Intriguingly, there is little overlap between the sets of genes linked to SATB2-dependent 3D epigenome effects at individual hierarchical levels. However, combining all level-specific gene sets, we find that SATB2 affects the spatial organization of almost half of the genes within neuron- and cognition-related GO categories. Ubiquitously expressed regulators of chromatin conformation like CTCF and cohesin appear to be instrumental in establishing or maintaining general aspects of 3D genome structure in all cell types^92^. Our analysis of chromosome conformation in *Satb2* cKO vs floxed cortical neurons indicates that in highly specialized cell types, such as pyramidal cells, additional layers of 3D genome organization co-exist that enable specific, adaptive transcriptional responses, tailored towards highly specialized biological processes^93,94^. These chromatin-architectural functions likely depend on proteins with highly restricted expression pattern, such as SATB2, which within the adult CNS is almost exclusively expressed in pyramidal neurons that form the cellular foundation of cognitive processes. Our data also indicate that the ubiquitous and the specialized layers of 3D genome organization may not necessarily operate independently. The enrichment of CTCF peaks at anchors of non-promoter based differential loops and the extensive overlap between SATB2 and CTCF interactors point towards potential cooperation between CTCF and SATB2 in establishing cell type-specific chromatin loops. Our data are consistent with a newly emerging literature, which indicates that CTCF co-interactors, in particular those that bind to its RNA-binding region^63^, and CTCF peak co-localizing factors^60^ modulate the strength of CTCF-mediated loops^95^. Prominent examples of CTCF co-localizing factors^60^ that also interact with SATB2 are BCLAF1 and NR2F2, two transcription factors highly expressed in pyramidal neurons, and HDAC1, a SATB2 interactor in upper layer cortical neurons^96^.

Extending previous findings that FIREs/super-FIREs are partially dependent on CTCF and cohesin^65^, we identify a highly selective subset of FIREs/super-FIREs that is SATB2-dependent. A hallmark of this subset is its enormous specificity since it harbors genes that are highly enriched (odds ratios over 10) for specialized cognition-related genes, i.e. the mouse orthologues of the genes discovered within GCA and NCS meta-loci in humans^67^. Neurodevelopmental gene sets in GCA-relevant meta-loci are expressed during prenatal-early childhood, whereas synaptic gene sets in NCS-relevant meta-loci are expressed postnatally. Our modelling of the spatial genome organization within the virtual 3D nucleus space provides a potential mechanism for their co-regulation. SATB2-dependent regulation of proximity in intra-nuclear positioning likely affects connectivity between cis-regulatory elements. Thereby, SATB2 might influence the formation of higher-order transcription hubs, which have recently emerged as a central mechanism, determining transcriptional bursting/co-activation of multiple genes within a functional gene set^97,98^.

*SATB2* has been identified as a locus conferring risk for SZ^99,100^ and associated with general cognitive ability^10^. Yet, thorough understanding of its contribution to these traits is currently missing. By leveraging common and rare genetic variation data, we uncover SATB2-dependent 3D epigenome remodeling as a candidate mechanism contributing specifically to human cognitive function and SZ. Lifting-over our data from mouse to human genome, we show that human orthologues of SATB2-dependent 3D genome sets are enriched for genes associated IQ and SZ. Even more surprisingly, human orthologues of non-coding regions undergoing SATB2-dependent chromatin modifications are enriched for SNP-based heritability for EA, IQ, and SZ. This indicates that the function of SATB2 in orchestrating expression of cognition-related genes, via coordinated 3D epigenome effects at multiple structural levels, has remained conserved during evolution of complex vertebrate brains. Furthermore, our study provides mechanistic insights and genetic support for the importance of SATB2-dependent mechanisms in human cognitive performance and links common and rare variants within SATB2 gene regulatory networks to the etiology of neuropsychiatric and neurodevelopmental diseases.

## Methods

### Primary cortical cultures

Neonatal Satb2^flx/flx^::Nes-Cre and Satb2^flx/flx^ mice^12^ were used to prepare primary cortical cultures, as previously described^16^. DIV 14 cortical neurons were first pre-treated with NBQX (20 μM, BioTechne) for 16 h, and then treated with bicucculine (50 μM, BioTechne) for 1 h. All experiments including animals were approved by the Austrian Animal Experimentation Ethics Board.

### Library preparation and sequencing

#### Hi-C

For *in situ* Hi-C, Arima-HiC kit was used, according to the manufacturer’s protocol. Briefly, 2.5×10^6^ primary cortical neurons were fixed directly on the dish by addition of formaldehyde to a final concentration of 1 % for 10 min on RT. After washing twice with 1X PBS cells were detached using a cell lifter and collected in 1X PBS containing protease inhibitors. Cells were pelleted at 500 g for 10 min at 4 °C and pellets were directly snap-frozen in liquid nitrogen.

After nuclei isolation and chromatin digestion, biotin was added and filled ends were ligated. After reverse crosslinking and purification, DNA was shared using a M220 Covaris sonicator at duty factor 12 %, PIP 75, CPB 200 for 60 s to achieve input fragment size of 400 bp. Additionally, DNA was size-selected using AMPure XP beads, according to Arima HiC kit manual. Hi-C libraries were prepared using Swift Biosciences Accel-NGS 2S Plus DNA Library Kit according to Arima-HiC Kit User Guide (November 2018).

#### ATAC-seq

10^6^ primary cortical neurons plated on a 35 mm dish were washed twice with 1X PBS before addition of nuclei isolation buffer (1X PBS, 0.1 % Triton X-100, 5 mM AEBSF Pefabloc, 5 mM Na-Butyrate, 1 tablet cOmplete EDTA-free Protease inhibitor). The dish was incubated on a shaker at 4°C for 7 min before detaching the cells with a cell lifter and gently resuspending them with a P1000 pipet. Cell suspension was transferred to an Eppendorf tube and spun down at 500 g at 4 °C for 10 min. Supernatant was removed and pellet was gently resuspended in 1X PBS with Protease inhibitors. Intact nuclei were counted in a Neubauer chamber. 25 K nuclei were used for tagmentation in 4X TD buffer (132 mM Tris-acetate pH 7.8, 264 mM Potassium-acetate, 40 mM Magnesium Acetate, 64 % Dimethylformamide, 0.005 % Digitonin and 1.5 μl Tn5 Transposase (15027865 Illumina)). Reaction was incubated at 37 °C for 30 min and purified using the QIAGEN MinElute Kit (28004 QIAGEN) according to the manufacturers protocol. Libraries were prepared as previously published^101^. Each library was given a unique barcode and amplified for 11 cycles. Afterwards, the PCR reaction was purified using 1X AMPURE XP Beads and eluted in 33 μl of 10 mM Tris HCl pH 8.0. Libraries were analyzed with Qubit and fragment size was accessed using the high sensitivity D-5000 screen tapes (5067-5592 Agilent) on a Tapestation.

#### CUT&RUN

After washing twice with 1X PBS, nuclei of 10^6^ cells plated on a 35 mm dish were isolated by addition of nuclei extraction buffer (NEB, 20 mM Hepes (KOH) pH 7.9, 10 mM KCl, 0.5 mM Spermidine, 0.1 % Triton X-100, 20 % Glycerol, 1 tablet cOmplete EDTA-free Protease inhibitor). Cells were incubated at 4 °C for 15 min, lifted from the dish using a cell lifter and gently resuspended using a P1000 pipet. Lysate was transferred to an Eppendorf tube and spun down at 600 g at 4 °C for 3 min. Pellet was resuspended in NEB and intact nuclei were counted in a Neubauer chamber. 100 K nuclei were used for each CUT&RUN reaction.

CUT&RUN protocol was adapted from the previously published protocol^102^. After Concavalin-A conjugated beads activation (21-1401-EPC Epicypher), nuclei were demobilized and blocked in blocking buffer (20 mM Hepes pH 7.5, 150 mM NaCl, 0.5 mM Spermidine, 0.1 % BSA, 2 mM EDTA, 1 tablet cOmplete EDTA-free Protease inhibitor) at RT for 5 min. After washing in washing buffer (20 mM Hepes pH 7.5, 150 mM NaCl, 0.5 mM Spermidine, 0.1 % BSA, 1 tablet cOmplete EDTA-free Protease inhibitor) primary antibody (Anti-SATB2 ab92446, Abcam) was added 1:100 in 250 μl of cold washing buffer and incubated at 4 °C for 2 h. Reaction was washed two additional times before pAG-MNAse (15-1016-EPC Epicypher) was added 1:20 in washing buffer and incubated for 1h at 4°C. Next, beads were washed twice before addition of 100 mM CaCl2 for 30 min at 0 °C in an ice water bath. 2X STOP buffer (200 mM NaCl, 20 mM EDTA, 4 mM EGTA, 50 μg/ml RNAseA, 40 μg/ml Glycogen, 10 pg/ml *E*.*Coli* spike-in DNA) was added and beads were incubated for 20 min at 37 °C to release DNA fragments and to digest RNA. DNA was purified by Phenol-Chloroform extraction and precipitated in 95 % EtOH at -80 °C. Pellet was dissolved in TE buffer pH 8.0 (12090015, ThermoFisher) and stored at -20 °C.

Library was prepared using the NEBNext Ultra Library prep Kit (E7645S, New England Biolabs) according to the manufacturer’s protocol. Adapters were used according to the DNA input at a working concentration of 1.5 μM. Library was PCR amplified with unique indexes (E7335S, New England Biolabs) for 14 cycles. After clean-up fragment size and concentration was analyzed before sending for sequencing.

All premade libraries were sent to Novogene UK for sequencing. After quality control, libraries were subjected to 150 bp paired-end sequencing (Illumina Novaseq platform).

### Data analysis

#### ATAC-seq

Pair-end data were mapped by using BWA-MEM (v0.7.17)^103^ and the resulting SAM files were converted into sorted and indexed BAM files (SAMtools v1.1)^104^. Reproducibility of replicate ATAC-seq libraries was assessed by calculating correlation between BAM files by using *deepTools2*^105^. Peaks were identified per experimental condition by employing Genrich tool (v0.6.1). Additionally, a consensus peak file across all conditions was produced by using *bedtools “merge”*. The number of read counts per region was calculated by using *featureCounts* (subread v.2.0.1)^106^. Next, differential analysis was done by using *edgeR*^107^. Principal component analysis showed a technical bias and samples were batch-normalized using RUVr method of RUVseq^108^. Homer (v4.11) was used for motif analysis. Genomic annotation and GO enrichment analysis was done by using *ChiPseeker*^29^ and *clusterProfiler*^109^. Footprinting analysis and occupancy prediction was performed by using *Tobias* (v0.12.10)^45^ over 5 kb-targets of promoter-based loops. Proximal Flx and cKO dOCRs were annotated using GREATs^59^ basal plus extention mode (Proximal: 5.0 kb upstream, 1.0 kb downstream, plus Distal: up to 2.5 kb).

#### CUT&RUN

After running a quality control on raw reads with FastQC, Illumina Universal adapters were trimmed with *Trim Galore!* with default parameters (--paired --illumina). Next, the trimmed reads were aligned to the mouse genome with *Bowtie2* (v2.4.5)^110^ with the following settings: bowtie2 --dovetail --local --very-sensitive-local --no-unal --no-mixed --no-discordant -q --phred33 -I 10 -X 700 -p 48 -x mm10. Subsequently, the mapped reads were filtered with *SAMtools* view. Reads with a mapping quality score (MAPQ) < 2 as well as optical and PCR duplicates are excluded. The reproducibility of replicate CUT&RUN libraries was assessed by calculating correlation between BAM files by using deepTools2^105^. Replicate BAM files were merged and handled as single file for the next steps. Genome coverage bedgraph files were computed with bamCoverage (bin size 50 bp) and normalized to reads per kilobase per million (RPKM) to minimize bias arising from different sequencing depths and batches. Unspecific noise was filtered out by subtracting the mean signal from each value of the corresponding bedgraph files. Peaks were called with *SEACR* v1.4^111^ under relaxed conditions and normalized to the IgG control bedgraph file. Genomic annotation and GO enrichment analysis was done by using *ChIPseeker*^29^ and *clusterProfiler*^109^. Peaks were annotated using *annotatr* package^112^ and defined as promoter-bound, if overlapping with gene promoters (1 kb upstream of TSS). Genomic enrichment analysis was performed by *Homer* (v4.11).

#### Hi-C

##### 1. Mapping, filtering and normalization

Paired-end sequencing reads were mapped to the mouse reference genome assembly (mm10), artifacts were filtered, and libraries was ICED normalized using the *HiC-Pro* (v.2.11.4)^113^. Data reproducibility was assessed using HiCRep method^25^ with a modified function renamed as ‘*HiCReproducibity*.*R’*. Briefly, Juicer Hi-C files were transformed into compatible Hi-C matrices at 250 kb resolution and observed frequencies per chromosome were extracted. Next, pairwise reproducibility values were computed using the function *get*.*scc* with the following parameters: *h=1, lbr = 0, ubr = 5000000*. Results were transformed into a quadratic matrix as input for the *pheatmap* R package^114^ for visualization purpose.

Contact distance decay plots were generated by using the function ‘*DecayFreq’
s* that computes and aggregates ICED normalized frequencies at 100 kb resolution.

##### 2. Compartment analysis

Saddle plots, visualizing genome compartmentalisation, were generated using *FAN-C* toolkit^46^. Compartment strength as defined by Flyamer et al., 2017 was calculated by taking natural logarithm of the AA * BB / AB^2.

Differential compartment analysis was carried out by using *dcHiC*^48^. Input file was created following *dcHiC* guidelines using HiC-Pro matrices at 100 kb.

To annotate genes residing in differential compartments, we used *annotatr* package^112^. For gene ontology analysis, we used only genes which promoter (defined as sequences 1-5 kb upstream of the TSS and < 1 kb upstream of the TSS) overlapped with differential compartments and had expression value base mean > 100 counts^16^.

##### 3. Topological associated domains (TADs)

TADs were computed at 25 kb resolution using *SpectralTAD*^52^. Briefly, HiC-Pro matrices were converted into bedpe format using the ‘*hicpro2bedpe’* function from *HiCcompare*^116^. Then, pooled samples were used as input for *SpectralTAD*, which was run iteratively for each chromosome.

Differential TAD boundaries between cKO and Flx Hi-C contact matrices were identified by *TADCompare*^52^. First, *SpectralTAD* TAD caller was used to pre-define TAD boundaries and then they were compared between the genotypes. Regions with absolute differential boundary score is > 2 (P value smaller than 0.05) were considered as differential boundaries. To annotate genes near differential boundaries, GREAT tool was used^59^, with the following association rule: basal + extension: 5 kb upstream, 1 kb downstream and 100 kb max extension.

##### 4. Chromatin loops

Chromatin loops were called by *Mustache* (v1.2.0), a local enrichment-based loop calling method^117^. Replicate Hi-C matrices were pooled, which allowed for conducting the analysis at 5 kb resolution. The following parameters for differential loop detection were used: P value (-p) of 0.05 and sparsity (-st) of 0.8 thresholds together with KR normalization. To assign differential chromatin loops to genes, we intersected loop anchors and targets with gene promoters (defined as sequences 1-5 kb upstream of the TSS and < 1 kb upstream of the TSS) using *annotatr*^112^. Loops annotated to genes overlapping between cKO and Flx dLoops were considered as shared promoter dLoops.

##### 5. Frequently interacting regions (FIREs)

Frequently interacting regions (FIREs) were called using the *FIREcaller* tool^66^. Briefly, ‘*sparseToDense*.*py’ HiC-Pr*o utility was used to create quadratic matrices at a resolution of 10 kb. To be able to run *FIREcaller*, a mappability file corresponding to *ArimaHiC* fragmentation was required. We provide this file for mm10 here: https://github.com/sespesogil/SATB2_3D_Genome/FIREs/resources. We provide code examples for FIRE calling (see “Code Availability”). Differential FIREs were identified as previously described^64^. Flx-specific FIRES were defined as genomic regions that have Flx FIRE scores greater than qnorm (0.975) and cKO FIRE scores lower than qnorm (0.9). Conversely, cKO-specific FIREs were defined as genomic regions with cKO FIRE score greater qnorm (0.975) and Flx FIRE score lower than qnorm (0.9). Flx- and cKO specific super-FIREs were identified by filtering out the super-FIREs common to both samples on genome coordinate level. To link differential FIREs and superFIREs to genes, *annotatr* package was used ^112^ to intersect FIREs and superFIREs with all genic annotations (1-5 kb upstream of the TSS, promoter (< 1 kb upstream of the TSS), 5UTR, first exons, exons, introns, CDS, 3UTR).

##### 6. 3D genome modelling

Hi-C data from floxed and Satb2 cKO cortical neurons were used to model chromatin conformation in 3D. TADs were called for each of the pooled HiC libraries (floxed and cKO) at 50 kb resolution and KR normalized using Arrowhead^118^. Next, we created a consensus TAD file for both conditions by merging the two bed files using bedtools merge and used it as an input file for *Chrom3D*^72^. We generated gtracks using https://github.com/sespesogil/automat_chrom3D. The analysis was restricted to diploid autosomal interactions at 50 kb and 1 Mb for intra and inter-chromosomal interactions, respectively. *Chrom3D* was run with the nucleus parameter set, with a radius of 3 μm for 1 million iterations: -r 3.0 -n 1000000 -l 5000 --nucleus. Mapping regions to Flx vs cKO 3D models was performed using: https://github.com/sespesogil/automat_chrom3D_colors. The coordinates of the promoters of mouse orthologues of NCS and GCA genes were obtained by using “AnnotationHub” package (v3.4.0). Pairwise Euclidean distances between regions of interest were computed using: https://github.com/sespesogil/automat_euclidean. The same utility was applied to compute hierarchical clustering using the complete linkage method (‘hclust’ function), integrated in *pheatmap R* package^114^. The resulting cluster dendograms were compared using *dendextend* package^119^ and the function ‘CutTheTree.R’ (https://github.com/sespesogil/automat_euclidean). The following parameters were computed: *entanglement* (measures the quality of the alignment of the two trees, varies between 1 (full entanglement) and 0 (no entanglement)), “Baker’s Gamma Index”, and “*Cophenetic*” correlation coefficients (both measuring similarity between tree topologies. Statistical significance of Baker’s Gamma Index was calculated by a permutation test with 1000 iterations. Models were visualized using ChimeraX (v1.0)^120^.

#### Gene Ontology enrichment analysis

*Metascape*^121^ was used to identify GO terms that were overrepresented in “SATB2 3D genome” sets such as genes residing in differential compartments, genes nearest to Flx and cKO TAD boundaries, dOCR-promoter interactions, Flx and cKO dLoop-promoter interactions, common promoter dLoop-promoter interactions, and genes residing in Flx and cKO FIREs/super-FIREs. As ontology sources, GO Molecular Functions, GO Biological Processes, and GO Cellular components were used. Terms with a P value < 0.01, a minimum count of 3, and an enrichment factor > 1.5 (the ratio between the observed counts and the counts expected by chance) were collected and grouped into clusters based on their membership similarities. P values were calculated based on the accumulative hypergeometric distribution, and q-values were calculated using the Banjamini-Hochberg procedure to account for multiple testing. All genes in the genome were used as enrichment background.

GO enrichment analysis of genes at SATB2 peaks and OCRs was performed by using *ChIPseeker*^29^ and *clusterProfiler*^109^.

Gene products associated with GO terms “axon development”, “postsynapse”, “neuron projection morphogenesis”, “glutamatergic synapse”, “cognition”, “T-cell activation”, “immune response” were downloaded from http://amigo.geneontology.org/amigo/landing. To test for significant overlap between “SATB2 3D genome” sets and genes associated with these GO categories, Fisher’s exact test was used. GREATv4 genes (GREATv4.genes.mm10.tsv) that were based on extremely high-confidence gene predictions were used as background set.

#### Overrepresentation analysis

We employed Fisher’s exact test to compare different gene lists with the genes engaged in SATB2-dependent distal regulatory element-promoter interactions (cKO dOCR-gene sets, Flx dLoop-gene set, common promoter dLoop-gene set), genes located within 100 kb regions flanking differential boundaries (Flx dBoundary gene set), genes mapped to differential compartments (dCompartment gene set), and genes mapped to Flx FIREs/super-FIREs (Flx FIRE/super-FIRE gene sets).

The list of DEGs between floxed and cKO neurons was obtained from Feurle et al.^16^, IEGs from Tyssowski et al.^34^, MEF2C-regulated genes from Harrington et al.^37^, and the genes, mapped to General Cognitive Ability (GCA) and Non-Cognitive Skills (NCS) meta-loci from Lam et al.^67^. GREATv4 genes (GREATv4.genes.mm10.tsv) that were based on extremely high-confidence gene predictions were used as background reference set (n = 21395).

#### Association analysis of genomic regions

A dedicated wrapper named ‘Enriched permutate’ was created to perform permutation analysis of genomic regions (see Code availability). Briefly, it computes the probability of genomic features to overlap using *RegionR* package^122^. Permutations were performed using bins of 5 kb (loop anchors) and 1 kb (SATB2 peaks and OCR) over mm10 genome, and 500 bp (lifted-over dOCR) over hg38 genome with a total number of 100000 iterations. For permutations of mouse regulatory elements, ChromHMM 18-state model of mouse forebrain (postnatal day 0) was used (ENCSR301UGN). Before permutations, multiple enhancer states (EnhG, Enh, EnhLo, EnhPois, EnhPr), Transcription Start Site states (Tss, TssFlnk, TssBiv), repressive states (ReprPC and ReprPCWk), quiescent states (QuiesG, Quies, Quies2, Quies3, and Quies4), and the two states associated with actively transcribed genes (Tx and TxWk) were first concatenated into a representative single state (Enh, Tss, ReprPC, Quies, Tx). For association analysis of human regulatory elements, ChromHMM 18-state model (ENCSR867UKF) of frontal cortex female adult (67 years) and female adult (80 years) from donor(s) ENCDO311AAA, ENCDO312AAA was used (ENCFF382TUC). Enhancer states (Enh A1, EnhA2, EnhBiv, EnhG1, EnhG2, and EnhWk) were concatenated as described above.

For enrichment analysis of histone marks and CTCF peaks, the following datasets were used (postnatal 0 day mouse forebrain): CTCF (ENCFF430PPJ); H3K27ac (ENCFF676TSV); H3K4me3 (ENCFF160SCR).

#### Conventional MAGMA and Hi-C-coupled MAGMA (H-MAGMA) gene-set analyses

Gene-set analysis (GSA) was used to analyze the effect of multiple SNPs and their joint effect toward a phenotype within defined genes of interest. GSA was performed using MAGMA^74^ using test and control GWAS summary statistics. First, SNPs from European 1000 Genomes phase 3 cohort (https://ctg.cncr.nl/software/MAGMA/ref_data/g1000_eur.zip) were annotated in relation to their location within genes on the GRCh37/hg19 human build using start/stop coordinates (https://ctg.cncr.nl/software/MAGMA/aux_files/NCBI37.3.zip) and a 20 kb window. Next, P values for each genes association with the relevant phenotypes were calculated, using a linear principal components regression model accounting for linkage disequilibrium (LD) between SNPs and employing 1000 Genomes European cohort data as a reference panel (gene-based analysis). Finally, a competitive GSA was performed based on P values from the gene-based analysis, testing if the gene-set has a stronger association with the phenotype of interest than other genes in the genome. MAGMA-GSA also accounted for gene size and gene density^123^.

While MAGMA uses locational proximity to annotate SNPs to genes, H-MAGMA incorporates tissue-specific Hi-C chromatin interactions to assign SNPs to genes^75^. The previously generated annotation files (Cortical_Neurons.genes.annot and Fetal_brain.genes.annot)^64^ based on cortical neuronal and fetal brain Hi-C data were used to provide SNP-to-gene relationships. Gene-based analysis and competitive GSA were performed as in MAGMA above.

#### GWAS Data

For both MAGMA and H-MAGMA analyses, gene sets were tested for enrichment of genes associated with five neurodevelopmental test phenotypes using genome-wide association study (GWAS) summary statistics for schizophrenia (SCZ)^49^, Intelligence (IQ)^10^, Educational Attainment (EA)^124^, Bipolar Disorder (BPD)^125^, and Major Depressive Disorder (MDD)^126^. In addition to the test phenotypes, five control phenotypes were also used: Alzheimers disease (AD)^127^, Stroke^128^, Coronary Artery Disease (CAD)^129^, Crohns Disease(CD)^130^, and Type 2 Diabetes (T2D)^131^.

#### Stratified Linkage Disequilibrium Score Regression (sLDSC)

To investigate if genomic regions were enriched for heritability contributing to neuropsychiatric disorders and cognition-associated phenotypes, stratified Linkage Disequilibrium Score Regression (sLDSC) (https://github.com/bulik/ldsc)^132,133^ was performed. Regions on mm10 mouse genome were lifted over to the human hg19 genome using UCSC liftOver (https://genome.ucsc.edu/cgi-bin/hgLiftOver) with default settings. To avoid significantly larger human regions inflating heritability enrichment values, lifted over regions more than 2 SD larger than the mean were removed. HapMap Project phase 3 SNPs with a MAF > 0.05 in the lifted over regions were considered in this analysis. LD scores between SNPs within a 1 centimorgan (cM) window were estimated using the 1000 Genomes Phase 3 European reference panel. SNP heritability for each phenotype was stratified within each set of regions using a model accounting for heritability associated with 53 functional genomic annotations found in the baseline model^132^. Enrichment for heritability compared to the baseline model was calculated with a corresponding P value. Enrichments surviving a Bonferroni-corrected P < 0.00125 were considered as significantly enriched.

#### Enrichment analysis for genes containing de novo mutations

Lists of *de novo* mutations (DNMs) identified in patients with ASD (n = 6,430), ID (n = 192) and in unaffected siblings (n = 1995) and controls (n = 54) based on exome sequencing of trios were sourced from^79^ and^80^. Genes containing DNMs reported in SZ patients (n = 3394) were taken from^77^ and^78^. DNMs identified in Developmental Disorder (DD) patients were sourced from^81^ but were subject to additional filtering based on posterior probability of *de novo* mutations, as described in^81^. DNMs were categorized as synonymous, missense and loss-of-function (includes nonsense, frameshift and splice site mutations). Each gene-set was tested for enrichments for genes harbouring DNMs in each phenotype using the R package *denovolyzeR*^134^. To do this, *denovolyzeR* counts the numbers of observed DNMs and derives the expected number of DNMs in a given population based on the mutability of the gene and the number of trios sequenced^134^. Enrichment of DNMs in a test gene-set was investigated using a two-sample Poisson rate ratio test, using the ratio of observed to expected DNMs in genes outside of the gene-set as a background model.

## Code availability

Study analysis, pipelines and code are available in the following Github repository: https://github.com/sespesogil/SATB2_3D_Genome.

## Datasets used in this study

GSE157375.

## Data availability

Raw data will be available as GEO record.

## Acknowledgements

This work was supported by Austrian Science Fund (FWF grants FWF-DK W1206 “Signal Processing in Neurons” to GD, FWF-SFB F44 “Cell Signaling in Chronic CNS Disorders” to GD and GA, FWF-P33027-B to GA, FWF-P32850-B to GD). We thank Drs. Isabella Cera and Patrick Feurle for their assistance in primary culture preparation.

## Notes

### Competing Interest Statement

The authors have declared no competing interest.

## References

1. Medrano-Fernández, A., and Barco, A. (2016). Nuclear organization and 3D chromatin architecture in cognition and neuropsychiatric disorders. Mol Brain 9, 83. 10.1186/s13041-016-0263-x.

2. Rajarajan, P., Gil, S.E., Brennand, K.J., and Akbarian, S. (2016). Spatial genome organization and cognition. Nat Rev Neurosci 17, 681–691. 10.1038/nrn.2016.124.

3. Bharadwaj, R., Peter, C.J., Jiang, Y., Roussos, P., Vogel-Ciernia, A., Shen, E.Y., Mitchell, A.C., Mao, W., Whittle, C., Dincer, A., et al. (2014). Conserved higher-order chromatin regulates NMDA receptor gene expression and cognition. Neuron 84, 997–1008. 10.1016/j.neuron.2014.10.032.

4. Bharadwaj, R., Jiang, Y., Mao, W., Jakovcevski, M., Dincer, A., Krueger, W., Garbett, K., Whittle, C., Tushir, J.S., Liu, J., et al. (2013). Conserved chromosome 2q31 conformations are associated with transcriptional regulation of GAD1 GABA synthesis enzyme and altered in prefrontal cortex of subjects with schizophrenia. J Neurosci 33, 11839–11851. 10.1523/JNEUROSCI.1252-13.2013.

5. Rajarajan, P., Borrman, T., Liao, W., Schrode, N., Flaherty, E., Casiño, C., Powell, S., Yashaswini, C., LaMarca, E.A., Kassim, B., et al. (2018). Neuron-specific signatures in the chromosomal connectome associated with schizophrenia risk. Science 362, eaat4311. 10.1126/science.aat4311.

6. Wang, D., Liu, S., Warrell, J., Won, H., Shi, X., Navarro, F.C.P., Clarke, D., Gu, M., Emani, P., Yang, Y.T., et al. (2018). Comprehensive functional genomic resource and integrative model for the human brain. Science 362, eaat8464. 10.1126/science.aat8464.

7. Zeisel, A., Hochgerner, H., Lönnerberg, P., Johnsson, A., Memic, F., van der Zwan, J., Häring, M., Braun, E., Borm, L.E., la Manno, G., et al. (2018). Molecular Architecture of the Mouse Nervous System. Cell 174, 999-1014.e22. 10.1016/J.CELL.2018.06.021.

8. Huang, Y., Song, N.-N., Lan, W., Hu, L., Su, C.-J., Ding, Y.-Q., and Zhang, L. (2013). Expression of transcription factor Satb2 in adult mouse brain. Anat Rec (Hoboken) 296, 452–461. 10.1002/ar.22656.

9. Szemes, M., Gyorgy, A., Paweletz, C., Dobi, A., and Agoston, D. v (2006). Isolation and characterization of SATB2, a novel AT-rich DNA binding protein expressed in development- and cell-specific manner in the rat brain. Neurochem Res 31, 237–246. 10.1007/s11064-005-9012-8.

10. Savage, J.E., Jansen, P.R., Stringer, S., Watanabe, K., Bryois, J., de Leeuw, C.A., Nagel, M., Awasthi, S., Barr, P.B., Coleman, J.R.I., et al. (2018). Genome-wide association meta-analysis in 269,867 individuals identifies new genetic and functional links to intelligence. Nat Genet 50, 912–919. 10.1038/s41588-018-0152-6.

11. Whitton, L., Apostolova, G., Rieder, D., Dechant, G., Rea, S., Donohoe, G., and Morris, D.W. (2018). Genes regulated by SATB2 during neurodevelopment contribute to schizophrenia and educational attainment. PLoS Genet 14, e1007515. 10.1371/journal.pgen.1007515.

12. Cera, I., Whitton, L., Donohoe, G., Morris, D.W., Dechant, G., and Apostolova, G. (2019). Genes encoding SATB2-interacting proteins in adult cerebral cortex contribute to human cognitive ability. PLoS Genet 15, e1007890. 10.1371/journal.pgen.1007890.

13. Zarate, Y.A., Smith-Hicks, C.L., Greene, C., Abbott, M.A., Siu, V.M., Calhoun, A.R.U.L., Pandya, A., Li, C., Sellars, E.A., Kaylor, J., et al. (2018). Natural history and genotype-phenotype correlations in 72 individuals with SATB2-associated syndrome. Am J Med Genet A 176, 925– 935. 10.1002/ajmg.a.38630.

14. Jaitner, C., Reddy, C., Abentung, A., Whittle, N., Rieder, D., Delekate, A., Korte, M., Jain, G., Fischer, A., Sananbenesi, F., et al. (2016). Satb2 determines miRNA expression and long-term memory in the adult central nervous system. Elife 5. 10.7554/elife.17361.

15. Li, Y., You, Q.L., Zhang, S.R., Huang, W.Y., Zou, W.J., Jie, W., Li, S.J., Liu, J.H., Lv, C.Y., Cong, J., et al. (2017). Satb2 Ablation Impairs Hippocampus-Based Long-Term Spatial Memory and Short-Term Working Memory and Immediate Early Genes (IEGs)-Mediated Hippocampal Synaptic Plasticity. Mol Neurobiol, 1–16. 10.1007/s12035-017-0531-5.

16. Feurle, P., Abentung, A., Cera, I., Wahl, N., Ablinger, C., Bucher, M., Stefan, E., Sprenger, S., Teis, D., Fischer, A., et al. (2021). SATB2-LEMD2 interaction links nuclear shape plasticity to regulation of cognition-related genes. EMBO J 40, e103701. 10.15252/embj.2019103701.

17. FitzPatrick, D.R., Carr, I.M., McLaren, L., Leek, J.P., Wightman, P., Williamson, K., Gautier, P., McGill, N., Hayward, C., Firth, H., et al. (2003). Identification of SATB2 as the cleft palate gene on 2q32-q33. Hum Mol Genet 12, 2491–2501. 10.1093/hmg/ddg248.

18. Wang, Z., Yang, X., Guo, S., Yang, Y., Su, X.-C., Shen, Y., and Long, J. (2014). Crystal structure of the ubiquitin-like domain-CUT repeat-like tandem of special AT-rich sequence binding protein 1 (SATB1) reveals a coordinating DNA-binding mechanism. J Biol Chem 289, 27376–27385. 10.1074/jbc.M114.562314.

19. Wang, Z., Yang, X., Chu, X., Zhang, J., Zhou, H., Shen, Y., and Long, J. (2012). The structural basis for the oligomerization of the N-terminal domain of SATB1. Nucleic Acids Res 40, 4193– 4202. 10.1093/nar/gkr1284.

20. Bell, R.A.V., Al-Khalaf, M.H., Brunette, S., Alsowaida, D., Chu, A., Bandukwala, H., Dechant, G., Apostolova, G., Dilworth, F.J., and Megeney, L.A. (2022). Chromatin Reorganization during Myoblast Differentiation Involves the Caspase-Dependent Removal of SATB2. Cells 2022, Vol. 11, Page 966 11, 966. 10.3390/CELLS11060966.

21. Pradhan, S.J., Reddy, P.C., Smutny, M., Sharma, A., Sako, K., Oak, M.S., Shah, R., Pal, M., Deshpande, O., Dsilva, G., et al. (2021). Satb2 acts as a gatekeeper for major developmental transitions during early vertebrate embryogenesis. Nature Communications 2021 12:1 12, 1– 19. 10.1038/s41467-021-26234-7.

22. Urrutia, G.A., Ramachandran, H., Cauchy, P., Boo, K., Ramamoorthy, S., Boller, S., Dogan, E., Clapes, T., Trompouki, E., Torres-Padilla, M.E., et al. (2021). ZFP451-mediated SUMOylation of SATB2 drives embryonic stem cell differentiation. Genes Dev 35, 1142–1160. 10.1101/GAD.345843.120/-/DC1.

23. Hardingham, G.E., Fukunaga, Y., and Bading, H. (2002). Extrasynaptic NMDARs oppose synaptic NMDARs by triggering CREB shut-off and cell death pathways. Nat Neurosci 5, 405– 414. 10.1038/nn835.

24. Peter J Skene, Jorja G Henikoff, S.H. (2018). Targeted in situ genome-wide profiling with high efficiency for low cell numbers. Nat Protoc 13, 1006–1019.

25. Yang, T., Zhang, F., Yardımci, G.G., Song, F., Hardison, R.C., Noble, W.S., Yue, F., and Li, Q. (2017). HiCRep: assessing the reproducibility of Hi-C data using a stratum-adjusted correlation coefficient. Genome Res 27, 1939–1949. 10.1101/GR.220640.117/-/DC1.

26. Espeso-Gil, S., Holik, A.Z., Bonnin, S., Jhanwar, S., Chandrasekaran, S., Pique-Regi, R., Albaigès-Ràfols, J., Maher, M., Permanyer, J., Irimia, M., et al. (2021). Environmental Enrichment Induces Epigenomic and Genome Organization Changes Relevant for Cognition. Front Mol Neurosci 14, 76. 10.3389/FNMOL.2021.664912/BIBTEX.

27. Davis, C.A., Hitz, B.C., Sloan, C.A., Chan, E.T., Davidson, J.M., Gabdank, I., Hilton, J.A., Jain, K., Baymuradov, U.K., Narayanan, A.K., et al. (2018). The Encyclopedia of DNA elements (ENCODE): data portal update. Nucleic Acids Res 46, D794–D801. 10.1093/NAR/GKX1081.

28. Ernst, J., and Kellis, M. (2017). Chromatin-state discovery and genome annotation with ChromHMM. Nat Protoc 12, 2478–2492. 10.1038/nprot.2017.124.

29. Yu, G., Wang, L.G., and He, Q.Y. (2015). ChIPseeker: an R/Bioconductor package for ChIP peak annotation, comparison and visualization. Bioinformatics 31, 2382–2383. 10.1093/BIOINFORMATICS/BTV145.

30. McKenna, W.L., Ortiz-Londono, C.F., Mathew, T.K., Hoang, K., Katzman, S., and Chen, B. (2015). Mutual regulation between Satb2 and Fezf2 promotes subcerebral projection neuron identity in the developing cerebral cortex. Proc Natl Acad Sci U S A 112, 11702–11707. 10.1073/pnas.1504144112.

31. Pradhan, S.J., Reddy, P.C., Smutny, M., Sharma, A., Sako, K., Oak, M.S., Shah, R., Pal, M., Deshpande, O., Tang, Y., et al. (2020). Satb2 acts as a gatekeeper for major developmental transitions during early vertebrate embryogenesis. bioRxiv, 2020.11.23.394171. 10.1101/2020.11.23.394171.

32. Urrutia, G.A., Ramachandran, H., Cauchy, P., Boo, K., Ramamoorthy, S., Boller, S., Dogan, E., Clapes, T., Trompouki, E., Torres-Padilla, M.E., et al. (2021). ZFP451-mediated SUMOylation of SATB2 drives embryonic stem cell differentiation. Genes Dev 35, 1142–1160. 10.1101/GAD.345843.120/-/DC1.

33. Guo, Q., Wang, Y., Wang, Q., Qian, Y., Jiang, Y., Dong, X., Chen, H., Chen, X., Liu, X., Yu, S., et al. (2022). In the developing cerebral cortex: axonogenesis, synapse formation, and synaptic plasticity are regulated by SATB2 target genes. Pediatric Research 2022, 1–9. 10.1038/s41390-022-02260-z.

34. Tyssowski, K.M., DeStefino, N.R., Cho, J.-H., Dunn, C.J., Poston, R.G., Carty, C.E., Jones, R.D., Chang, S.M., Romeo, P., Wurzelmann, M.K., et al. (2018). Different Neuronal Activity Patterns Induce Different Gene Expression Programs. Neuron 98, 530-546.e11. 10.1016/J.NEURON.2018.04.001.

35. Yap, E.-L., and Greenberg, M.E. (2018). Activity-Regulated Transcription: Bridging the Gap between Neural Activity and Behavior. Neuron 100, 330–348. 10.1016/J.NEURON.2018.10.013.

36. Chen, L.F., Zhou, A.S., and West, A.E. (2017). Transcribing the connectome: Roles for transcription factors and chromatin regulators in activity-dependent synapse development. J Neurophysiol 118, 755–770. 10.1152/JN.00067.2017/ASSET/IMAGES/LARGE/Z9K0071742110003.JPEG.

37. Harrington, A.J., Raissi, A., Rajkovich, K., Berto, S., Kumar, J., Molinaro, G., Raduazzo, J., Guo, Y., Loerwald, K., Konopka, G., et al. (2016). MEF2C regulates cortical inhibitory and excitatory synapses and behaviors relevant to neurodevelopmental disorders. Elife 5. 10.7554/ELIFE.20059.

38. Flavell, S.W., Kim, T.K., Gray, J.M., Harmin, D.A., Hemberg, M., Hong, E.J., Markenscoff-Papadimitriou, E., Bear, D.M., and Greenberg, M.E. (2008). Genome-Wide Analysis of MEF2 Transcriptional Program Reveals Synaptic Target Genes and Neuronal Activity-Dependent Polyadenylation Site Selection. Neuron 60, 1022–1038. 10.1016/J.NEURON.2008.11.029.

39. Bedogni, F., and Hevner, R.F. (2021). Cell-Type-Specific Gene Expression in Developing Mouse Neocortex: Intermediate Progenitors Implicated in Axon Development. Front Mol Neurosci 14, 132. 10.3389/FNMOL.2021.686034/BIBTEX.

40. Rodriguez, M., Choi, J., Park, S., and Sockanathan, S. (2012). Gde2 regulates cortical neuronal identity by controlling the timing of cortical progenitor differentiation. Development 139, 3870–3879. 10.1242/DEV.081083.

41. Swayne, L.A., and Bennett, S.A.L. (2016). Connexins and pannexins in neuronal development and adult neurogenesis. BMC Cell Biol 17, 39–49. 10.1186/S12860-016-0089-5/FIGURES/3.

42. van der Velde, A., Fan, K., Tsuji, J., Moore, J.E., Purcaro, M.J., Pratt, H.E., and Weng, Z. (2021). Annotation of chromatin states in 66 complete mouse epigenomes during development. Communications Biology 2021 4:1 4, 1–15. 10.1038/s42003-021-01756-4.

43. Dunham, I., Kundaje, A., Aldred, S.F., Collins, P.J., Davis, C.A., Doyle, F., Epstein, C.B., Frietze, S., Harrow, J., Kaul, R., et al. (2012). An integrated encyclopedia of DNA elements in the human genome. Nature 2012 489:7414 489, 57–74. 10.1038/nature11247.

44. Lu, L., Liu, X., Huang, W.K., Giusti-Rodríguez, P., Cui, J., Zhang, S., Xu, W., Wen, Z., Ma, S., Rosen, J.D., et al. (2020). Robust Hi-C Maps of Enhancer-Promoter Interactions Reveal the Function of Non-coding Genome in Neural Development and Diseases. Mol Cell 79, 521-534.e15. 10.1016/j.molcel.2020.06.007.

45. Bentsen, M., Goymann, P., Schultheis, H., Klee, K., Petrova, A., Wiegandt, R., Fust, A., Preussner, J., Kuenne, C., Braun, T., et al. (2020). ATAC-seq footprinting unravels kinetics of transcription factor binding during zygotic genome activation. Nature Communications 2020 11:1 11, 1–11. 10.1038/s41467-020-18035-1.

46. Kruse, K., Hug, C.B., and Vaquerizas, J.M. (2020). FAN-C: a feature-rich framework for the analysis and visualisation of chromosome conformation capture data. Genome Biol 21, 303. 10.1186/s13059-020-02215-9.

47. Wutz, G., Várnai, C., Nagasaka, K., Cisneros, D.A., Stocsits, R.R., Tang, W., Schoenfelder, S., Jessberger, G., Muhar, M., Hossain, M.J., et al. (2017). Topologically associating domains and chromatin loops depend on cohesin and are regulated by CTCF, WAPL, and PDS5 proteins. EMBO J 36, 3573–3599. 10.15252/EMBJ.201798004.

48. Chakraborty, A., Wang, J., and Ay, F. (2022). dcHiC: differential compartment analysis of Hi-C datasets. bioRxiv, 2021.02.02.429297. 10.1101/2021.02.02.429297.

49. Trubetskoy, V., Pardiñas, A.F., Qi, T., Panagiotaropoulou, G., Awasthi, S., Bigdeli, T.B., Bryois, J., Chen, C.Y., Dennison, C.A., Hall, L.S., et al. (2022). Mapping genomic loci implicates genes and synaptic biology in schizophrenia. Nature 2022 604:7906 604, 502–508. 10.1038/s41586-022-04434-5.

50. Lam, M., Hill, W.D., Trampush, J.W., Yu, J., Knowles, E., Davies, G., Stahl, E., Huckins, L., Liewald, D.C., Djurovic, S., et al. (2019). Pleiotropic Meta-Analysis of Cognition, Education, and Schizophrenia Differentiates Roles of Early Neurodevelopmental and Adult Synaptic Pathways. The American Journal of Human Genetics 105, 334–350. 10.1016/J.AJHG.2019.06.012.

51. Cresswell, K.G., Stansfield, J.C., and Dozmorov, M.G. (2020). SpectralTAD: An R package for defining a hierarchy of topologically associated domains using spectral clustering. BMC Bioinformatics 21, 1–19. 10.1186/S12859-020-03652-W/FIGURES/4.

52. Cresswell, K.G., and Dozmorov, M.G. (2020). TADCompare: An R Package for Differential and Temporal Analysis of Topologically Associated Domains. Front Genet 11, 158. 10.3389/FGENE.2020.00158/BIBTEX.

53. Dixon, J.R., Selvaraj, S., Yue, F., Kim, A., Li, Y., Shen, Y., Hu, M., Liu, J.S., and Ren, B. (2012). Topological domains in mammalian genomes identified by analysis of chromatin interactions. Nature 2012 485:7398 485, 376–380. 10.1038/nature11082.

54. Dixon, J.R., Jung, I., Selvaraj, S., Shen, Y., Antosiewicz-Bourget, J.E., Lee, A.Y., Ye, Z., Kim, A., Rajagopal, N., Xie, W., et al. (2015). Chromatin architecture reorganization during stem cell differentiation. Nature 2015 518:7539 518, 331–336. 10.1038/nature14222.

55. Lazar, N.H., Nevonen, K.A., O’Connell, B., McCann, C., O’Neill, R.J., Green, R.E., Meyer, T.J., Okhovat, M., and Carbone, L. (2018). Epigenetic maintenance of topological domains in the highly rearranged gibbon genome. Genome Res 28, 983–997. 10.1101/GR.233874.117/-/DC1.

56. Winick-Ng, W., Kukalev, A., Harabula, I., Zea-Redondo, L., Szabó, D., Meijer, M., Serebreni, L., Zhang, Y., Bianco, S., Chiariello, A.M., et al. (2021). Cell-type specialization is encoded by specific chromatin topologies. Nature 2021 599:7886 599, 684–691. 10.1038/s41586-021-04081-2.

57. Jerković, I., and Cavalli, G. (2021). Understanding 3D genome organization by multidisciplinary methods. Nature Reviews Molecular Cell Biology 2021 22:8 22, 511–528. 10.1038/s41580-021-00362-w.

58. Ong, C.-T., and Corces, V.G. (2014). CTCF: an architectural protein bridging genome topology and function. Nat Rev Genet 15, 234–246. 10.1038/nrg3663.

59. McLean, C.Y., Bristor, D., Hiller, M., Clarke, S.L., Schaar, B.T., Lowe, C.B., Wenger, A.M., and Bejerano, G. (2010). GREAT improves functional interpretation of cis-regulatory regions. Nat Biotechnol 28, 495. 10.1038/NBT.1630.

60. Hu, G., Dong, X., Gong, S., Song, Y., Hutchins, A.P., and Yao, H. (2020). Systematic screening of CTCF binding partners identifies that BHLHE40 regulates CTCF genome-wide distribution and long-range chromatin interactions. Nucleic Acids Res 48, 9606–9620. 10.1093/NAR/GKAA705.

61. Oughtred, R., Rust, J., Chang, C., Breitkreutz, B.J., Stark, C., Willems, A., Boucher, L., Leung, G., Kolas, N., Zhang, F., et al. (2021). The BioGRID database: A comprehensive biomedical resource of curated protein, genetic, and chemical interactions. Protein Science 30, 187–200. 10.1002/PRO.3978.

62. Marcon, E., Ni, Z., Pu, S., Turinsky, A.L., Trimble, S.S., Olsen, J.B., Silverman-Gavrila, R., Silverman-Gavrila, L., Phanse, S., Guo, H., et al. (2014). Human-Chromatin-Related Protein Interactions Identify a Demethylase Complex Required for Chromosome Segregation. Cell Rep 8, 297–310. 10.1016/J.CELREP.2014.05.050.

63. Saldaña-Meyer, R., Rodriguez-Hernaez, J., Escobar, T., Nishana, M., Jácome-López, K., Nora, E.P., Bruneau, B.G., Tsirigos, A., Furlan-Magaril, M., Skok, J., et al. (2019). RNA Interactions Are Essential for CTCF-Mediated Genome Organization. Mol Cell 76, 412-422.e5. 10.1016/j.molcel.2019.08.015.

64. Hu, B., Won, H., Mah, W., Park, R.B., Kassim, B., Spiess, K., Kozlenkov, A., Crowley, C.A., Pochareddy, S., Ashley-Koch, A.E., et al. (2021). Neuronal and glial 3D chromatin architecture informs the cellular etiology of brain disorders. Nature Communications 2021 12:1 12, 1–13. 10.1038/s41467-021-24243-0.

65. Schmitt, A.D., Hu, M., Jung, I., Xu, Z., Qiu, Y., Tan, C.L., Li, Y., Lin, S., Lin, Y., Barr, C.L., et al. (2016). A Compendium of Chromatin Contact Maps Reveals Spatially Active Regions in the Human Genome. Cell Rep 17, 2042–2059. 10.1016/j.celrep.2016.10.061.

66. Crowley, C., Yang, Y., Qiu, Y., Hu, B., Abnousi, A., Lipiński, J., Plewczyński, D., Wu, D., Won, H., Ren, B., et al. (2021). FIREcaller: Detecting frequently interacting regions from Hi-C data. Comput Struct Biotechnol J 19, 355–362. 10.1016/j.csbj.2020.12.026.

67. Lam, M., Chen, C.-Y., Hill, W.D., Xia, C., Tian, R., Levey, D.F., Gelernter, J., Stein, M.B., team, B.B., Hatoum, A.S., et al. (2021). Dissecting Biological Pathways of Psychopathology using Cognitive Genomics. medRxiv, 2021.07.20.21260487. 10.1101/2021.07.20.21260487.

68. Banerjee-Basu, S., and Packer, A. (2010). SFARI Gene: an evolving database for the autism research community. Dis Model Mech 3, 133–135. 10.1242/DMM.005439.

69. Gonzalez-Mantilla, A.J., Moreno-De-Luca, A., Ledbetter, D.H., and Martin, C.L. (2016). A Cross-Disorder Method to Identify Novel Candidate Genes for Developmental Brain Disorders. JAMA Psychiatry 73, 275–283. 10.1001/JAMAPSYCHIATRY.2015.2692.

70. Kochinke, K., Zweier, C., Nijhof, B., Fenckova, M., Cizek, P., Honti, F., Keerthikumar, S., Oortveld, M.A.W., Kleefstra, T., Kramer, J.M., et al. (2016). Systematic Phenomics Analysis Deconvolutes Genes Mutated in Intellectual Disability into Biologically Coherent Modules. The American Journal of Human Genetics 98, 149–164. 10.1016/J.AJHG.2015.11.024.

71. Stadhouders, R., Filion, G.J., and Graf, T. (2019). Transcription factors and 3D genome conformation in cell-fate decisions. Nature 2019 569:7756 569, 345–354. 10.1038/s41586-019-1182-7.

72. Paulsen, J., Sekelja, M., Oldenburg, A.R., Barateau, A., Briand, N., Delbarre, E., Shah, A., Sørensen, A.L., Vigouroux, C., Buendia, B., et al. (2017). Chrom3D: three-dimensional genome modeling from Hi-C and nuclear lamin-genome contacts. Genome Biol 18, 21. 10.1186/s13059-016-1146-2.

73. Espeso-Gil, S., Halene, T., Bendl, J., Kassim, B., ben Hutta, G., Iskhakova, M., Shokrian, N., Auluck, P., Javidfar, B., Rajarajan, P., et al. (2020). A chromosomal connectome for psychiatric and metabolic risk variants in adult dopaminergic neurons. Genome Med 12, 1–19. 10.1186/s13073-020-0715-x.

74. de Leeuw, C.A., Mooij, J.M., Heskes, T., and Posthuma, D. (2015). MAGMA: Generalized Gene-Set Analysis of GWAS Data. PLoS Comput Biol 11, e1004219. 10.1371/journal.pcbi.1004219.

75. Sey, N.Y.A., Hu, B., Mah, W., Fauni, H., McAfee, J.C., Rajarajan, P., Brennand, K.J., Akbarian, S., and Won, H. (2020). A computational tool (H-MAGMA) for improved prediction of brain-disorder risk genes by incorporating brain chromatin interaction profiles. Nature Neuroscience 2020 23:4 23, 583–593. 10.1038/s41593-020-0603-0.

76. Won, H., de la Torre-Ubieta, L., Stein, J.L., Parikshak, N.N., Huang, J., Opland, C.K., Gandal, M.J., Sutton, G.J., Hormozdiari, F., Lu, D., et al. (2016). Chromosome conformation elucidates regulatory relationships in developing human brain. Nature 538, 523–527. 10.1038/nature19847.

77. Howrigan, D.P., Rose, S.A., Samocha, K.E., Fromer, M., Cerrato, F., Chen, W.J., Churchhouse, C., Chambert, K., Chandler, S.D., Daly, M.J., et al. (2020). Exome sequencing in schizophrenia-affected parent–offspring trios reveals risk conferred by protein-coding de novo mutations. Nature Neuroscience 2020 23:2 23, 185–193. 10.1038/s41593-019-0564-3.

78. Rees, E., Han, J., Morgan, J., Carrera, N., Escott-Price, V., Pocklington, A.J., Duffield, M., Hall, L.S., Legge, S.E., Pardiñas, A.F., et al. (2020). De novo mutations identified by exome sequencing implicate rare missense variants in SLC6A1 in schizophrenia. Nature Neuroscience 2020 23:2 23, 179–184. 10.1038/s41593-019-0565-2.

79. Satterstrom, F.K., Kosmicki, J.A., Wang, J., Breen, M.S., de Rubeis, S., An, J.Y., Peng, M., Collins, R., Grove, J., Klei, L., et al. (2020). Large-Scale Exome Sequencing Study Implicates Both Developmental and Functional Changes in the Neurobiology of Autism. Cell 180, 568-584.e23. 10.1016/j.cell.2019.12.036.

80. Genovese, G., Fromer, M., Stahl, E.A., Ruderfer, D.M., Chambert, K., Landén, M., Moran, J.L., Purcell, S.M., Sklar, P., Sullivan, P.F., et al. (2016). Increased burden of ultra-rare protein-altering variants among 4,877 individuals with schizophrenia. Nat Neurosci 19, 1433–1441. 10.1038/nn.4402.

81. McRae, J.F., Clayton, S., Fitzgerald, T.W., Kaplanis, J., Prigmore, E., Rajan, D., Sifrim, A., Aitken, S., Akawi, N., Alvi, M., et al. (2017). Prevalence and architecture of de novo mutations in developmental disorders. Nature 2017 542:7642 542, 433–438. 10.1038/nature21062.

82. Bengani, H., Handley, M., Alvi, M., Ibitoye, R., Lees, M., Lynch, S.A., Lam, W., Fannemel, M., Nordgren, A., Malmgren, H., et al. (2017). Clinical and molecular consequences of disease-associated de novo mutations in SATB2. Genetics in Medicine 19, 900–908. 10.1038/gim.2016.211.

83. Dunham, I., Kundaje, A., Aldred, S.F., Collins, P.J., Davis, C.A., Doyle, F., Epstein, C.B., Frietze, S., Harrow, J., Kaul, R., et al. (2012). An integrated encyclopedia of DNA elements in the human genome. Nature 2012 489:7414 489, 57–74. 10.1038/nature11247.

84. Zhang, J.-P., Lencz, T., Geisler, S., DeRosse, P., Bromet, E.J., and Malhotra, A.K. (2013). Genetic variation in BDNF is associated with antipsychotic treatment resistance in patients with schizophrenia. Schizophr Res 146, 285–288. 10.1016/j.schres.2013.01.020.

85. Marco, A., Meharena, H.S., Dileep, V., Raju, R.M., Davila-Velderrain, J., Zhang, A.L., Adaikkan, C., Young, J.Z., Gao, F., Kellis, M., et al. (2020). Mapping the epigenomic and transcriptomic interplay during memory formation and recall in the hippocampal engram ensemble. Nat Neurosci. 10.1038/s41593-020-00717-0.

86. Fernandez-Albert, J., Lipinski, M., Lopez-Cascales, M.T., Rowley, M.J., Martin-Gonzalez, A.M., del Blanco, B., Corces, V.G., and Barco, A. (2019). Immediate and deferred epigenomic signatures of in vivo neuronal activation in mouse hippocampus. Nat Neurosci 22, 1718–1730. 10.1038/s41593-019-0476-2.

87. Baranek, C., Dittrich, M., Parthasarathy, S., Bonnon, C.G., Britanova, O., Lanshakov, D., Boukhtouche, F., Sommer, J.E., Colmenares, C., Tarabykin, V., et al. (2012). Protooncogene Ski cooperates with the chromatin-remodeling factor Satb2 in specifying callosal neurons. Proceedings of the National Academy of Sciences 109, 3546–3551. 10.1073/pnas.1108718109.

88. Britanova, O., de Juan Romero, C., Cheung, A., Kwan, K.Y., Schwark, M., Gyorgy, A., Vogel, T., Akopov, S., Mitkovski, M., Agoston, D., et al. (2008). Satb2 is a postmitotic determinant for upper-layer neuron specification in the neocortex. Neuron 57, 378–392. 10.1016/j.neuron.2007.12.028.

89. Nichols, M.H., and Corces, V.G. (2021). Principles of 3D compartmentalization of the human genome. Cell Rep 35, 109330. 10.1016/J.CELREP.2021.109330.

90. Rowley, M.J., and Corces, V.G. (2018). Organizational principles of 3D genome architecture. Nature Reviews Genetics 2018 19:12 19, 789–800. 10.1038/s41576-018-0060-8.

91. Nora, E.P., Goloborodko, A., Valton, A.L., Gibcus, J.H., Uebersohn, A., Abdennur, N., Dekker, J., Mirny, L.A., and Bruneau, B.G. (2017). Targeted Degradation of CTCF Decouples Local Insulation of Chromosome Domains from Genomic Compartmentalization. Cell 169, 930-944.e22. 10.1016/J.CELL.2017.05.004.

92. Merkenschlager, M., and Nora, E.P. (2016). CTCF and Cohesin in Genome Folding and Transcriptional Gene Regulation. http://dx.doi.org/10.1146/annurev-genom-083115-022339 17, 17–43. 10.1146/ANNUREV-GENOM-083115-022339.

93. Feng, D., Chen, Y., Dai, R., Bian, S., Xue, W., Zhu, Y., Li, Z., Yang, Y., Zhang, Y., Zhang, J., et al. (2022). Chromatin organizer SATB1 controls the cell identity of CD4+ CD8+ double-positive thymocytes by regulating the activity of super-enhancers. Nature Communications 2022 13:1 13, 1–17. 10.1038/s41467-022-33333-6.

94. Zelenka, T., Klonizakis, A., Tsoukatou, D., Franzenburg, S., Tzerpos, P., Papamatheakis, D.-A., Tzonevrakis, I.-R., Nikolaou, C., Plewczynski, D., and Spilianakis, C. (2021). The 3D enhancer network of the developing T cell genome is controlled by SATB1. bioRxiv, 2021.07.09.451769. 10.1101/2021.07.09.451769.

95. Hansen, A.S., Hsieh, T.H.S., Cattoglio, C., Pustova, I., Saldaña-Meyer, R., Reinberg, D., Darzacq, X., and Tjian, R. (2019). Distinct Classes of Chromatin Loops Revealed by Deletion of an RNA-Binding Region in CTCF. Mol Cell 76, 395-411.e13. 10.1016/J.MOLCEL.2019.07.039.

96. Gyorgy, A.B., Szemes, M., de Juan Romero, C., Tarabykin, V., and Agoston, D. v (2008). SATB2 interacts with chromatin-remodeling molecules in differentiating cortical neurons. Eur J Neurosci 27, 865–873. 10.1111/j.1460-9568.2008.06061.x.

97. Kim, S., and Shendure, J. (2019). Mechanisms of Interplay between Transcription Factors and the 3D Genome. Mol Cell 76, 306–319. 10.1016/J.MOLCEL.2019.08.010.

98. Lim, B., and Levine, M.S. (2021). Enhancer-promoter communication: hubs or loops? Curr Opin Genet Dev 67, 5–9. 10.1016/J.GDE.2020.10.001.

99. Giusti-Rodríguez, P., Lu, L., Yang, Y., Crowley, C.A., Liu, X., Juric, I., Martin, J.S., Abnousi, A., Allred, S.C., Ancalade, N., et al. (2019). Using three-dimensional regulatory chromatin interactions from adult and fetal cortex to interpret genetic results for psychiatric disorders and cognitive traits. bioRxiv 18, 406330. 10.1101/406330.

100. Ripke, S., Neale, B.M., Corvin, A., Walters, J.T.R., Farh, K.-H., Holmans, P.A., Lee, P., Bulik-Sullivan, B., Collier, D.A., Huang, H., et al. (2014). Biological insights from 108 schizophrenia-associated genetic loci. Nature 511, 421–427. 10.1038/nature13595.

101. Buenrostro, J.D., Wu, B., Chang, H.Y., and Greenleaf, W.J. (2015). ATAC-seq: A Method for Assaying Chromatin Accessibility Genome-Wide. Current protocols in molecular biology / edited by Frederick M. Ausubel … [et al.] 109, 21.29.1. 10.1002/0471142727.MB2129S109.

102. Hainer, S.J., Bošković, A., McCannell, K.N., Rando, O.J., and Fazzio, T.G. (2019). Profiling of Pluripotency Factors in Single Cells and Early Embryos. Cell 177, 1319-1329.e11. 10.1016/j.cell.2019.03.014.

103. Li, H. (2013). Aligning sequence reads, clone sequences and assembly contigs with BWA-MEM. 10.48550/arxiv.1303.3997.

104. Danecek, P., Bonfield, J.K., Liddle, J., Marshall, J., Ohan, V., Pollard, M.O., Whitwham, A., Keane, T., McCarthy, S.A., Davies, R.M., et al. (2021). Twelve years of SAMtools and BCFtools. Gigascience 10, 1–4. 10.1093/GIGASCIENCE/GIAB008.

105. Ramírez, F., Ryan, D.P., Grüning, B., Bhardwaj, V., Kilpert, F., Richter, A.S., Heyne, S., Dündar, F., and Manke, T. (2016). deepTools2: a next generation web server for deep-sequencing data analysis. Nucleic Acids Res 44, W160–W165. 10.1093/NAR/GKW257.

106. Liao, Y., Smyth, G.K., and Shi, W. (2014). featureCounts: an efficient general purpose program for assigning sequence reads to genomic features. Bioinformatics 30, 923–930. 10.1093/bioinformatics/btt656.

107. Robinson, M.D., McCarthy, D.J., and Smyth, G.K. (2010). edgeR: a Bioconductor package for differential expression analysis of digital gene expression data. Bioinformatics 26, 139–140. 10.1093/BIOINFORMATICS/BTP616.

108. Risso, D., Ngai, J., Speed, T.P., and Dudoit, S. (2014). Normalization of RNA-seq data using factor analysis of control genes or samples. Nat Biotechnol 32, 896–902. 10.1038/nbt.2931.

109. Wu, T., Hu, E., Xu, S., Chen, M., Guo, P., Dai, Z., Feng, T., Zhou, L., Tang, W., Zhan, L., et al. (2021). clusterProfiler 4.0: A universal enrichment tool for interpreting omics data. The Innovation 2, 100141. 10.1016/J.XINN.2021.100141.

110. Langmead, B., and Salzberg, S.L. (2012). Fast gapped-read alignment with Bowtie 2. Nature Methods 2012 9:4 9, 357–359. 10.1038/nmeth.1923.

111. Meers, M.P., Tenenbaum, D., and Henikoff, S. (2019). Peak calling by Sparse Enrichment Analysis for CUT&RUN chromatin profiling. Epigenetics Chromatin 12, 1–11. 10.1186/S13072-019-0287-4/FIGURES/6.

112. Cavalcante, R.G., and Sartor, M.A. (2017). annotatr: genomic regions in context. Bioinformatics 33, 2381–2383. 10.1093/BIOINFORMATICS/BTX183.

113. Servant, N., Varoquaux, N., Lajoie, B.R., Viara, E., Chen, C.J., Vert, J.P., Heard, E., Dekker, J., and Barillot, E. (2015). HiC-Pro: An optimized and flexible pipeline for Hi-C data processing. Genome Biol 16, 1–11. 10.1186/S13059-015-0831-X/TABLES/4.

114. Kolde, R., and others (2012). Pheatmap: pretty heatmaps. R package version 1, 726.

115. Flyamer, I.M., Gassler, J., Imakaev, M., Brandão, H.B., Ulianov, S. v., Abdennur, N., Razin, S. v., Mirny, L.A., and Tachibana-Konwalski, K. (2017). Single-nucleus Hi-C reveals unique chromatin reorganization at oocyte-to-zygote transition. Nature 2017 544:7648 544, 110–114. 10.1038/nature21711.

116. Stansfield, J.C., Cresswell, K.G., Vladimirov, V.I., and Dozmorov, M.G. (2018). HiCcompare: An R-package for joint normalization and comparison of HI-C datasets. BMC Bioinformatics 19, 1–10. 10.1186/S12859-018-2288-X/TABLES/3.

117. Roayaei Ardakany, A., Gezer, H.T., Lonardi, S., and Ay, F. (2020). Mustache: Multi-scale detection of chromatin loops from Hi-C and Micro-C maps using scale-space representation. Genome Biol 21, 1–17. 10.1186/S13059-020-02167-0/FIGURES/7.

118. Rao, S.S.P., Huntley, M.H., Durand, N.C., Stamenova, E.K., Bochkov, I.D., Robinson, J.T., Sanborn, A.L., Machol, I., Omer, A.D., Lander, E.S., et al. (2014). A 3D map of the human genome at kilobase resolution reveals principles of chromatin looping. Cell 159, 1665–1680. 10.1016/J.CELL.2014.11.021.

119. Galili, T. (2015). dendextend: an R package for visualizing, adjusting and comparing trees of hierarchical clustering. Bioinformatics 31, 3718–3720. 10.1093/BIOINFORMATICS/BTV428.

120. Goddard, T.D., Huang, C.C., Meng, E.C., Pettersen, E.F., Couch, G.S., Morris, J.H., and Ferrin, T.E. (2018). UCSF ChimeraX: Meeting modern challenges in visualization and analysis. Protein Science 27, 14–25. 10.1002/PRO.3235.

121. Zhou, Y., Zhou, B., Pache, L., Chang, M., Khodabakhshi, A.H., Tanaseichuk, O., Benner, C., and Chanda, S.K. (2019). Metascape provides a biologist-oriented resource for the analysis of systems-level datasets. Nature Communications 2019 10:1 10, 1–10. 10.1038/s41467-019-09234-6.

122. Gel, B., Díez-Villanueva, A., Serra, E., Buschbeck, M., Peinado, M.A., and Malinverni, R. (2016). regioneR: an R/Bioconductor package for the association analysis of genomic regions based on permutation tests. Bioinformatics 32, 289–291. 10.1093/BIOINFORMATICS/BTV562.

123. de Leeuw, C.A., Neale, B.M., Heskes, T., and Posthuma, D. (2016). The statistical properties of gene-set analysis. Nat Rev Genet 17, 353–364. 10.1038/NRG.2016.29.

124. Lee, J.J., Wedow, R., Okbay, A., Kong, E., Maghzian, O., Zacher, M., Nguyen-Viet, T.A., Bowers, P., Sidorenko, J., Karlsson Linnér, R., et al. (2018). Gene discovery and polygenic prediction from a genome-wide association study of educational attainment in 1.1 million individuals. Nat Genet 50, 1112–1121. 10.1038/s41588-018-0147-3.

125. Mullins, N., Forstner, A.J., O’Connell, K.S., Coombes, B., Coleman, J.R.I., Qiao, Z., Als, T.D., Bigdeli, T.B., Børte, S., Bryois, J., et al. (2021). Genome-wide association study of more than 40,000 bipolar disorder cases provides new insights into the underlying biology. Nature Genetics 2021 53:6 53, 817–829. 10.1038/s41588-021-00857-4.

126. Howard, D.M., Adams, M.J., Clarke, T.K., Hafferty, J.D., Gibson, J., Shirali, M., Coleman, J.R.I., Hagenaars, S.P., Ward, J., Wigmore, E.M., et al. (2019). Genome-wide meta-analysis of depression identifies 102 independent variants and highlights the importance of the prefrontal brain regions. Nature Neuroscience 2019 22:3 22, 343–352. 10.1038/s41593-018-0326-7.

127. Jansen, I.E., Savage, J.E., Watanabe, K., Bryois, J., Williams, D.M., Steinberg, S., Sealock, J., Karlsson, I.K., Hägg, S., Athanasiu, L., et al. (2019). Genome-wide meta-analysis identifies new loci and functional pathways influencing Alzheimer’s disease risk. Nature Genetics 2019 51:3 51, 404–413. 10.1038/s41588-018-0311-9.

128. Traylor, M., Farrall, M., Holliday, E.G., Sudlow, C., Hopewell, J.C., Cheng, Y.-C., Fornage, M., Ikram, M.A., Malik, R., Bevan, S., et al. (2012). Genetic risk factors for ischaemic stroke and its subtypes (the METASTROKE collaboration): a meta-analysis of genome-wide association studies. Lancet Neurol 11, 951–962. 10.1016/S1474-4422(12)70234-X.

129. Schunkert, H., König, I.R., Kathiresan, S., Reilly, M.P., Assimes, T.L., Holm, H., Preuss, M., Stewart, A.F.R., Barbalic, M., Gieger, C., et al. (2011). Large-scale association analysis identifies 13 new susceptibility loci for coronary artery disease. Nat Genet 43, 333–338. 10.1038/ng.784.

130. Liu, J.Z., van Sommeren, S., Huang, H., Ng, S.C., Alberts, R., Takahashi, A., Ripke, S., Lee, J.C., Jostins, L., Shah, T., et al. (2015). Association analyses identify 38 susceptibility loci for inflammatory bowel disease and highlight shared genetic risk across populations. Nat Genet 47, 979–986. 10.1038/ng.3359.

131. Mahajan, A., Taliun, D., Thurner, M., Robertson, N.R., Torres, J.M., Rayner, N.W., Payne, A.J., Steinthorsdottir, V., Scott, R.A., Grarup, N., et al. (2018). Fine-mapping type 2 diabetes loci to single-variant resolution using high-density imputation and islet-specific epigenome maps. Nature Genetics 2018 50:11 50, 1505–1513. 10.1038/s41588-018-0241-6.

132. Finucane, H.K., Bulik-Sullivan, B., Gusev, A., Trynka, G., Reshef, Y., Loh, P.-R., Anttila, V., Xu, H., Zang, C., Farh, K., et al. (2015). Partitioning heritability by functional annotation using genome-wide association summary statistics. Nat Genet 47, 1228–1235. 10.1038/ng.3404.

133. Bulik-Sullivan, B., Loh, P.R., Finucane, H.K., Ripke, S., Yang, J., Patterson, N., Daly, M.J., Price, A.L., Neale, B.M., Corvin, A., et al. (2015). LD Score regression distinguishes confounding from polygenicity in genome-wide association studies. Nature Genetics 2015 47:3 47, 291–295. 10.1038/ng.3211.

134. Ware, J.S., Samocha, K.E., Homsy, J., and Daly, M.J. (2015). Interpreting de novo Variation in Human Disease Using denovolyzeR. Curr Protoc Hum Genet 87, 7.25.1-7.25.15. 10.1002/0471142905.hg0725s87.

