## Supplemental Figures for "SATB2 organizes the 3D genome architecture of cognition in cortical neurons"

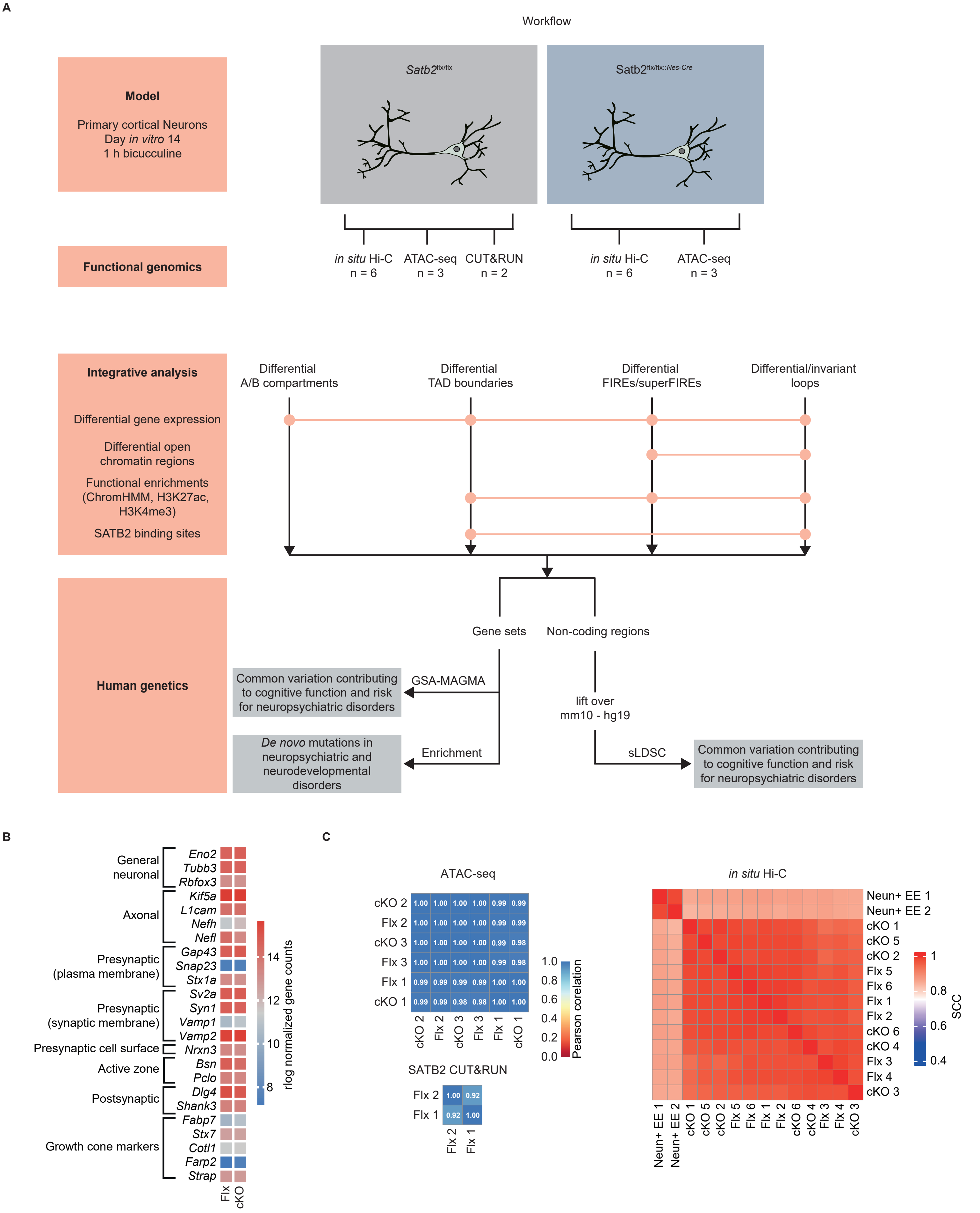
A
Workflow
Model
Primary cortical Neurons
Day in vitro 14
1 h bicucculine
Satb2flx/flx
Satb2flx/flx::Nes-Cre
Functional genomics
in situ Hi-C n = 6
ATAC-seq n = 3
CUT&RUN n = 2
in situ Hi-C n = 6
ATAC-seq n = 3
Integrative analysis
Differential gene expression
Differential open chromatin regions
Functional enrichments (ChromHMM, H3K27ac, H3K4me3)
SATB2 binding sites
Differential A/B compartments
Differential TAD boundaries
Differential FIREs/superFIREs
Differential/invariant loops
Human genetics
Gene sets
Non-coding regions
GSA-MAGMA
Common variation contributing to cognitive function and risk for neuropsychiatric disorders
Enrichment
De novo mutations in neuropsychiatric and neurodevelopmental disorders
lift over mm10 - hg19
sLDSC
Common variation contributing to cognitive function and risk for neuropsychiatric disorders
B
General neuronal
Eno2
Tubb3
Rbfox3
Axonal
Kif5a
L1cam
Nefh
Nefl
Presynaptic (plasma membrane)
Gap43
Snap23
Stx1a
Presynaptic (synaptic membrane)
Sv2a
Syn1
Vamp1
Vamp2
Presynaptic cell surface
Nrxn3
Active zone
Bsn
Pclo
Postsynaptic
Dlg4
Shank3
Growth cone markers
Fabp7
Stx7
Cotl1
Farp2
Strap
Flx
cKO
rlog normalized gene counts
14
12
10
8
C
ATAC-seq
cKO 2: 1.00 1.00 1.00 1.00 0.99 0.99
Flx 2: 1.00 1.00 1.00 1.00 0.99 0.99
cKO 3: 1.00 1.00 1.00 1.00 0.99 0.98
Flx 3: 1.00 1.00 1.00 1.00 0.99 0.98
Flx 1: 0.99 0.99 0.99 0.99 1.00 1.00
cKO 1: 0.99 0.99 0.98 0.98 1.00 1.00
cKO 2 Flx 2 cKO 3 Flx 3 Flx 1 cKO 1
Pearson coreltation
1.0
0.8
0.6
0.4
0.2
0.0
SATB2 CUT&RUN
Flx 2: 1.00 0.92
Flx 1: 0.92 1.00
Flx 2 Flx 1
in situ Hi-C
Neun+ EE 1
Neun+ EE 2
cKO 1
cKO 5
cKO 2
Flx 5
Flx 6
Flx 1
Flx 2
cKO 6
cKO 4
Flx 3
Flx 4
cKO 3
SCC
1
0.8
0.6
0.4

**Figure S1: Experimental and integrative analysis workflow, developmental and synaptic maturation markers in floxed vs cKO cultures, and quality control of ATAC-seq, Hi-C, and CUT&RUN libraries**

A. ATAC-seq and Hi-C libraries were generated from primary cultures derived from Satb2$^{flx/flx}$ and *Satb2*$^{flx/flx::\textit{Nes-Cre}}$ mice. CUT&RUN libraries were prepared from *Satb2*$^{flx/flx}$ cultures. Genotype-specific 3D genome architectural units such as compartments, TAD boundaries, FIREs, and chromatin loops were integrated with epigenomic and transcriptomic datasets. Human orthologues of the gene-sets derived from the differential analyses as well as mouse-to-human lifted-over genomic regions were used to analyze the effect of neuropsychiatric and neurodevelopmental disorder genetic risk.
B. Heatmap of neuronal marker expression values in floxed vs cKO cortical neurons, transcriptomic data from Feurle et al.[16] .
C. Pearson correlation coefficients of read counts between ATAC-seq and CUT&RUN library library replicates.
D. Stratum-adjusted correlation coefficients (SSC) analyzed by HiCRep[25] across replicate Hi-C libraries from this study and adult mouse cortex Hi-C dataset[26], EE, environmental enrichment.

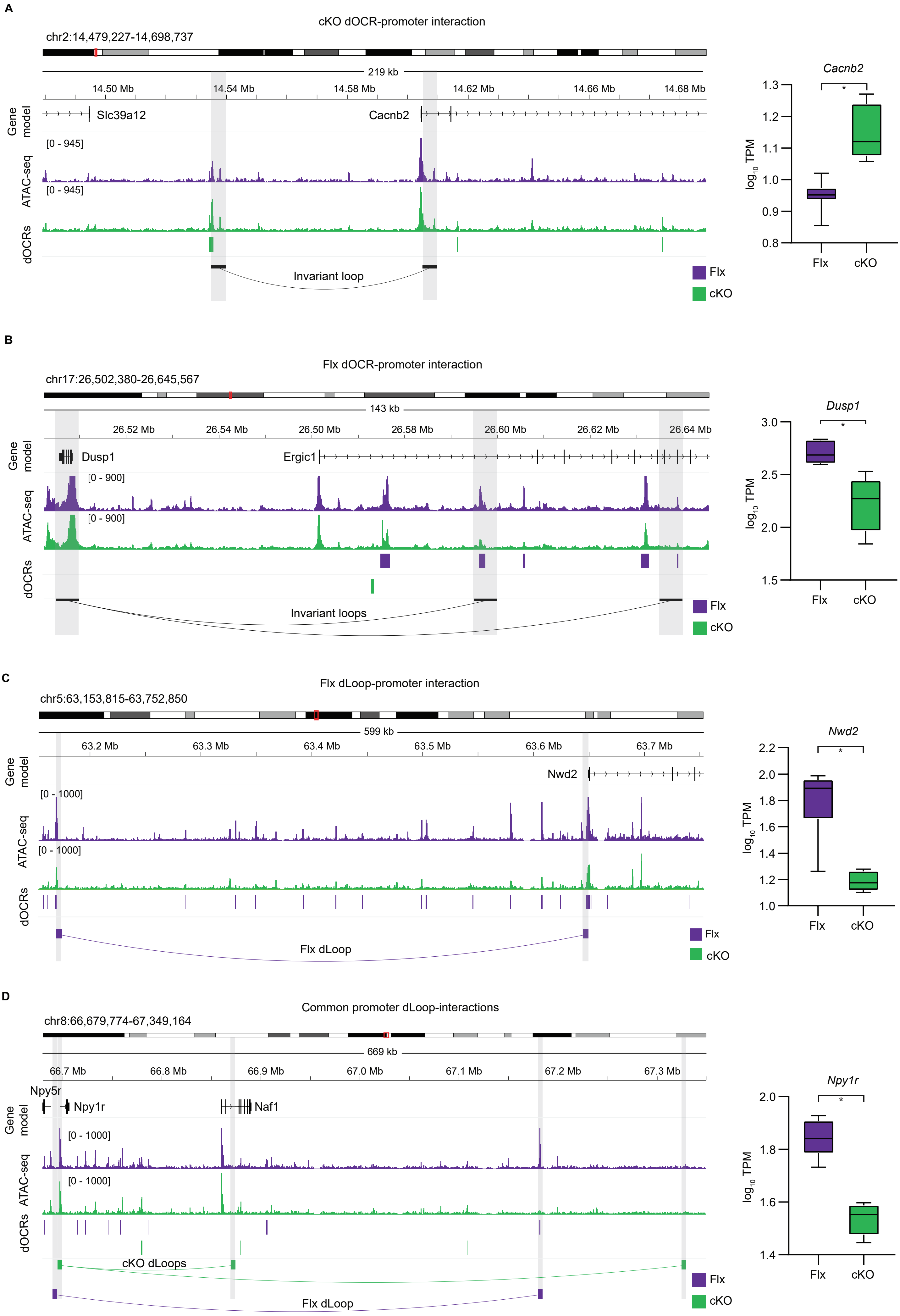
A
cKO dOCR-promoter interaction
chr2:14,479,227-14,698,737
219 kb
14.50 Mb
14.54 Mb
14.58 Mb
14.62 Mb
14.66 Mb
14.68 Mb
Gene model
Slc39a12
Cacnb2
ATAC-seq
[0 - 945]
[0 - 945]
dOCRs
Invariant loop
Flx
cKO
Cacnb2
log10 TPM
1.3
1.2
1.1
1.0
0.9
0.8
*
Flx
cKO
B
Flx dOCR-promoter interaction
chr17:26,502,380-26,645,567
143 kb
26.52 Mb
26.54 Mb
26.50 Mb
26.58 Mb
26.60 Mb
26.62 Mb
26.64 Mb
Gene model
Dusp1
Ergic1
ATAC-seq
[0 - 900]
[0 - 900]
dOCRs
Invariant loops
Flx
cKO
Dusp1
log10 TPM
3.0
2.5
2.0
1.5
*
Flx
cKO
C
Flx dLoop-promoter interaction
chr5:63,153,815-63,752,850
599 kb
63.2 Mb
63.3 Mb
63.4 Mb
63.5 Mb
63.6 Mb
63.7 Mb
Gene model
Nwd2
ATAC-seq
[0 - 1000]
[0 - 1000]
dOCRs
Flx dLoop
Flx
cKO
Nwd2
log10 TPM
2.2
2.0
1.8
1.6
1.4
1.2
1.0
*
Flx
cKO
D
Common promoter dLoop-interactions
chr8:66,679,774-67,349,164
669 kb
66.7 Mb
66.8 Mb
66.9 Mb
67.0 Mb
67.1 Mb
67.2 Mb
67.3 Mb
Gene model
Npy5r
Npy1r
Naf1
ATAC-seq
[0 - 1000]
[0 - 1000]
dOCRs
cKO dLoops
Flx dLoop
Flx
cKO
Npy1r
log10 TPM
2.0
1.8
1.6
1.4
*
Flx
cKO

**Figure S2: Representative examples of SATB2-dependent regulatory element-promoter interactions**

A. Representative example of cKO dOCR-promoter interaction. Left, IGV genome browser tracks showing chromatin accessibility in floxed and cKO neurons (colored tracks), dOCRs, and invariant chromatin loops (presented via black arcs). The anchors of an invariant loop, connecting *Cacnb2* promoter to cKO dOCR (region with increased accessibility in cKO vs floxed neurons), are highlighted grey. Right, boxplot showing expression levels of *Cacnb2* in floxed (n = 7) and cKO cultures (n = 7), respectively. TPM, Transcripts Per Million mapped reads, Padj = 2.01e-11, calculated by DESeq2 (two-sided Wald test), transcriptomic data from Feurle et al.[16].

B. Representative example of Flx dOCR-promoter interaction. IGV genome browser tracks showing chromatin accessibility in floxed and cKO neurons (colored tracks), dOCRs, and invariant chromatin loops (presented via black arcs). The anchors of two invariant loops, connecting *Dusp1* promoter to two Flx dOCRs (regions with increased accessibility in floxed vs cKO neurons), are highlighted grey. Right, boxplot showing expression levels of *Dusp1* in floxed (n = 7) and cKO cultures (n = 7), respectively. TPM, Transcripts Per Million mapped reads, P adj = 1.68e-14, calculated by DESeq2 (two-sided Wald test), transcriptomic data from Feurle et al.[16].

C. Representative example of Flx dLoop-promoter interaction. IGV genome browser tracks showing chromatin accessibility in floxed and cKO neurons (colored tracks), dOCRs, and dLoops (presented via colored arcs). The anchors of a Flx dLoop, connecting *Nwd2* promoter to a distal regulatory element, are highlighted grey. Right, boxplot showing expression levels of *Nwd2* in floxed (n = 7) and cKO cultures (n = 7), respectively. TPM, Transcripts Per Million mapped reads, P adj = 1.36e-28, calculated by DESeq2 (two-sided Wald test), transcriptomic data from Feurle et al[16].

D. Representative example of common promoter dLoop-interactions. Left, IGV genome browser tracks showing chromatin accessibility in floxed and cKO neurons (colored tracks), dOCRs, and dLoops (presented via colored arcs). The anchors of Flx and cKO dLoops, connecting *Npy1r* promoter to different distal regulatory elements in floxed vs cKO neurons, are highlighted grey. Right, boxplot showing expression levels of *Npy1r* in floxed (n = 7) and cKO cultures (n = 7), respectively. TPM, Transcripts Per Million mapped reads, P adj = 7.83e-36, calculated by DESeq2 (two-sided Wald test), transcriptomic data from Feurle et al.[16].

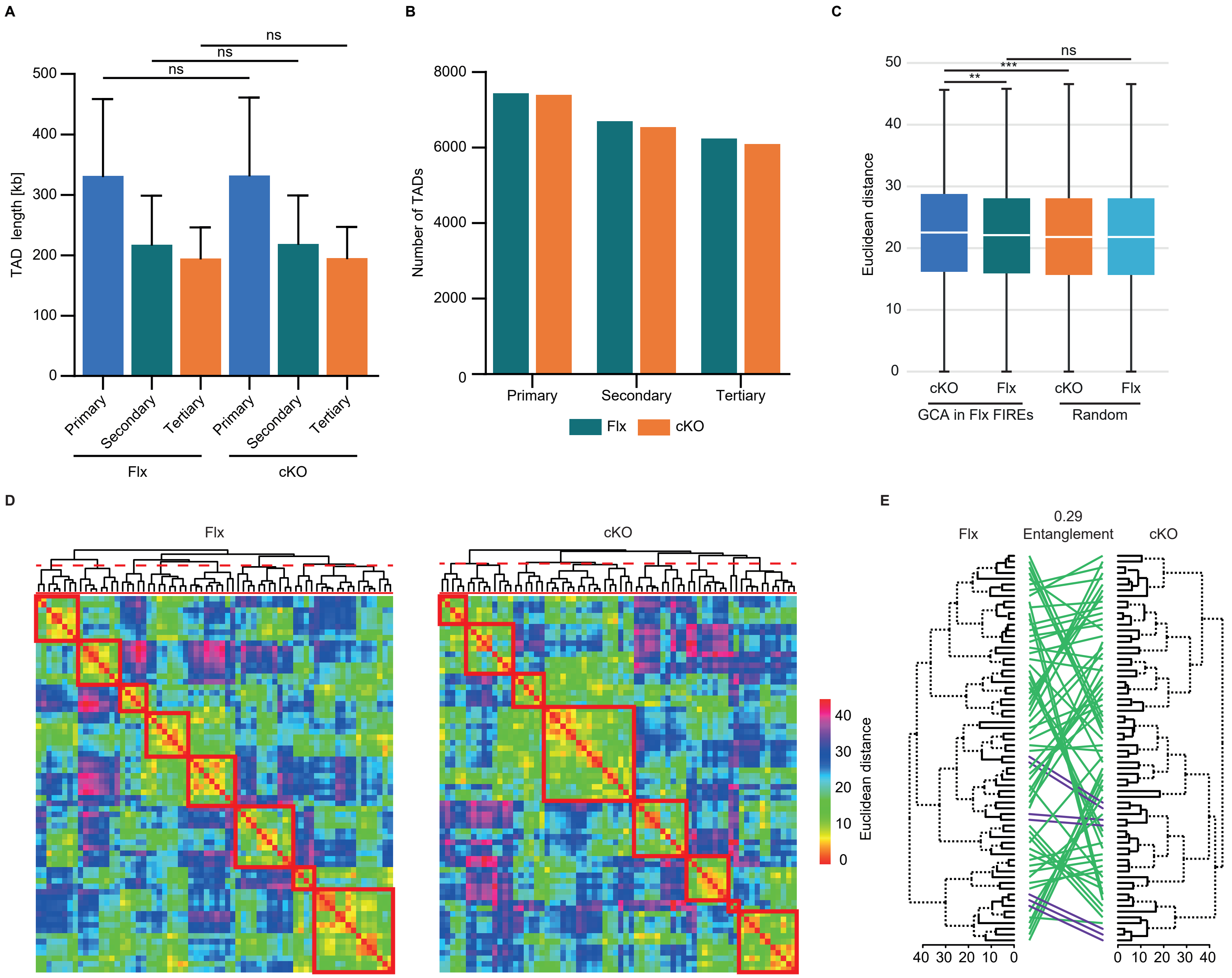
A
B
C
ns
ns
ns
TAD length [kb]
500
400
300
200
100
0
Primary
Secondary
Tertiary
Primary
Secondary
Tertiary
Flx
cKO
8000
6000
4000
2000
0
Number of TADs
Primary
Secondary
Tertiary
Flx
cKO
50
40
30
20
10
0
Euclidean distance
ns
***
**
cKO
Flx
cKO
Flx
GCA in Flx FIREs
Random
D
Flx
cKO
40
30
20
10
0
Euclidean distance
E
0.29
Entanglement
Flx
cKO
40 30 20 10 0
0 10 20 30 40

**Figure S3: Characterization of TADs in floxed vs cKO Hi-C matrices and Chrom3D genome modeling**

A. Length of TADs and sub-TADs (secondary and tertiary) called by SpectralTAD[50] in floxed vs cKO samples.
B. Number of called TADs and sub-TADs (secondary and tertiary) genotypes in floxed vs cKO samples.
C. Box plots showing the pairwise Euclidean distances (ED) between the 69 domains harboring Flx FIREs/super-FIRE genes with human orthologues mapped to GCA meta-loci in Flx vs cKO 3D genome models (P = 0.0097, Wilcoxon signed rank test). Shown are the pairwise ED between randomly permuted sets of domains of the same size (n = 1000 permutations) in Flx vs cKO 3D genome models (P = 0.32 GCA in Flx FIREs vs random domains, Flx 3D model; P = 4.29e-06, GCA in Flx FIREs vs random domains, cKO 3D model, Wilcoxon rank-sum tests).
D. Heatmaps of pairwise ED between the 69 domains harboring Flx FIREs/super-FIRE genes with human orthologues mapped to NCS meta-loci in Flx vs cKO 3D genome models. Red squares depict the Euclidean hot spots (clusters of domains that are spatially close together) in Flx and cKO sample.
E. Comparison between Flx and cKO dendograms of the 69 domains harboring Flx FIREs/super-FIRE genes with human orthologues mapped to GCA meta-loci. “Unique” nodes, with a combination of labels/items not present in the other tree, are highlighted with dashed lines. Green colored lines connect domains that belong to different subtrees in the two dendogrames.
